# Reference genome choice compromises population genetic analyses

**DOI:** 10.1101/2024.11.26.625554

**Authors:** Maria Akopyan, Matthew Genchev, Ellie E. Armstrong, Jazlyn A. Mooney

## Abstract

Characterizing genetic variation in natural populations is vital to evolutionary biology, however many non-model species lack genomic resources. Here, we demonstrate that reference bias significantly affects population genomic analyses by mapping whole genome sequence data from gray foxes (*Urocyon cinereoargenteus*) to a conspecific reference and two heterospecific canid genomes (dog and Arctic fox). Mapping to the conspecific genome improved read pairing by ∼5%, detected 26–32% more SNPs, and 33–35% more singletons. Nucleotide diversity estimates increased over 30%, *F*_ST_ increased from 0.189 to 0.197, and effective population size estimates were 30-60% higher with the conspecific reference. Recombination rates varied by up to 3-fold at chromosome ends with heterospecific references. Importantly, *F*_ST_ outlier detection differed markedly, with heterospecific genomes identifying twice as many unique outlier windows. These findings highlight the impact of reference genome choice and the importance of conspecific genomic resources for accurate evolutionary inference.

**Graphical Abstract:** 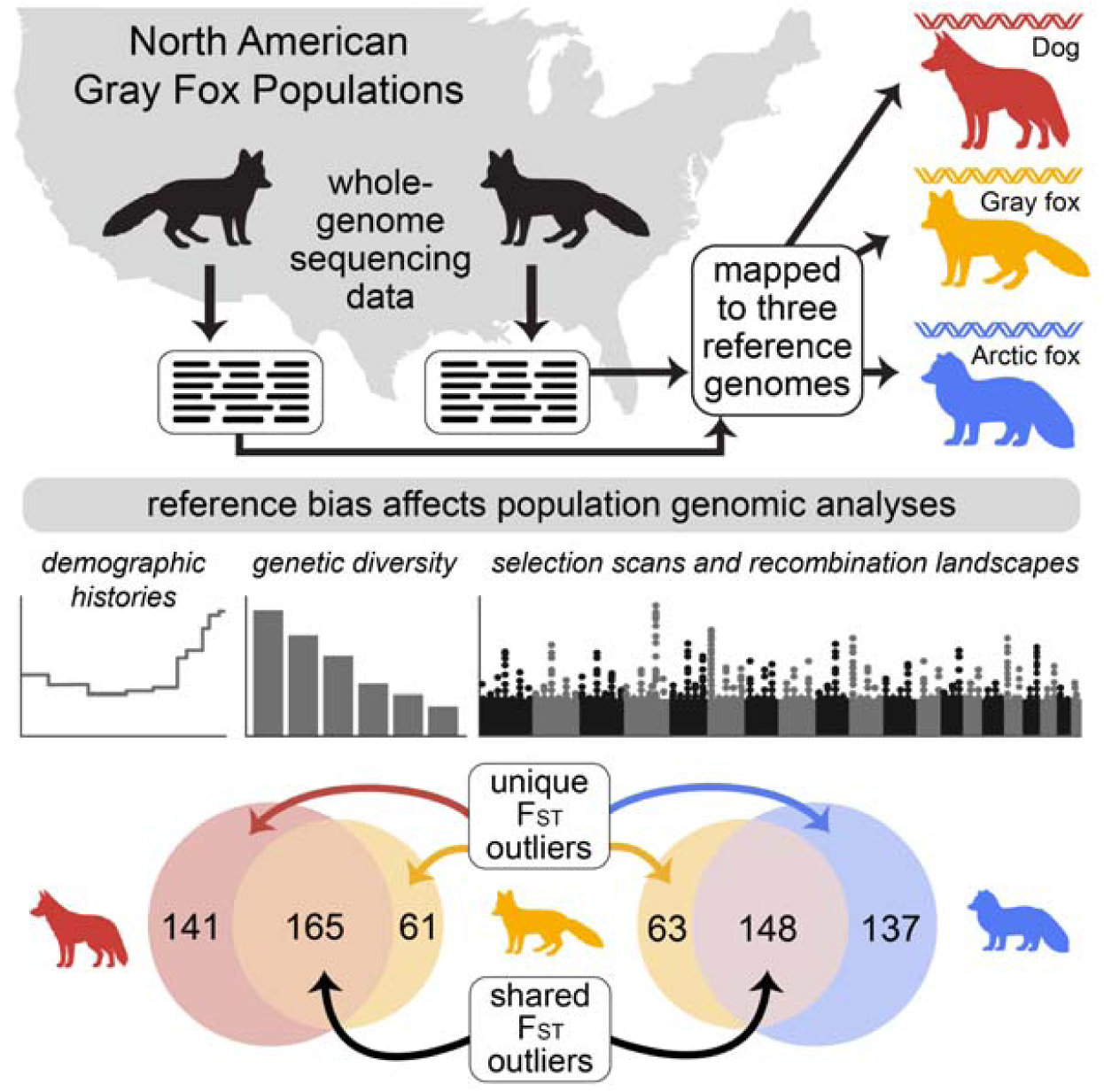

**Highlights:**

- A species-specific reference genome improves read mapping and variant detection
- Reference bias underestimates genetic diversity and differentiation
- Divergent reference genomes distort demographic histories and recombination landscapes
- Unique F_ST_ outliers are detected across references, affecting functional interpretations

## Introduction

Accurately characterizing genetic variation is essential for understanding the evolutionary dynamics of populations and informing management strategies for species of conservation concern. The increasing accessibility of whole genome sequencing (WGS) data for non-model organisms has led to a proliferation of studies employing WGS data that estimate genetic diversity, assess population structure and connectivity, and identify adaptive genetic variation. Recent advancements in analytical techniques for WGS data have enabled sophisticated analyses that extend beyond classic population genetic diversity metrics to characterize structural variation^1^, infer demographic histories^2^, and estimate population recombination rates^3^. These developments provide deeper insights into the historical and contemporary processes shaping genetic variation in natural populations. However, the reliability of these inferences and cross-population comparisons depends on the reference genome used. Reference genomes are essential for comparing individuals and populations, but they do not capture the full spectrum of genetic diversity. Population-level diversity is missed because a haploid reference represents variation from only a single individual at each site in the genome.

Reference bias arises because a single reference genome—usually a haploid sequence from one individual—is used as the coordinate system for mapping. This reference cannot fully represent the genetic diversity of an entire species. As a result, sequencing reads that closely match the reference tend to map with higher quality and are retained, while dissimilar reads with lower quality scores often map poorly and are discarded. This can lead to mapping errors or missed variant calls, especially for true population-specific genetic variation. Thus, reference bias will result in missed or mis-called genotypes that will impact estimates of genetic diversity and downstream analyses with the potential to distort conclusions about the population’s evolutionary history. Reference bias, along with sequencing depth, has been shown to influence downstream genetic analyses. For example, in simulated low-coverage (2-4x) genomes, introducing a 2% reference divergence can cause estimates of θ to increase by 0.14- to 0.18- fold^4^. Similar effects were noted on commonly used neutrality statistics: a small downward bias in Tajima’s D, an upward bias in Fu and Li’s D, and a strong downward bias in Fay and Wu’s H^4^. Importantly, these shifts were still observed even at moderate coverage (8x) and in empirical data^4^.

It is often the case that researchers will map to a reference genome from related species when a conspecific reference genome is unavailable. For example, in modern dogs and wolves, using a heterospecific (wolf) reference instead of a conspecific (dog) reference led to a 10% difference in heterozygosity estimates between individual dogs^5^. In non-model species like big cats, using a conspecific reference can alter both heterozygosity (ranging from a 0.25-fold decrease to a 3.21-fold increase) and demographic history inference^6^. In fishes, reference bias has been associated with overestimated heterozygosity, with local reference genomes reducing reference/non-reference heterozygous calls by 0.13-0.31% and non-reference/non-reference calls by 0.05-0.10%^7^. Studies have also shown that reference bias leads to inaccurate detection of structural variation^8^ and distorted phylogenies^9^, potentially skewing conclusions about population history and genetic diversity in fish species. Lastly, recent work on the impact of reference bias in two species of conservation concern, the New Zealand kiwi and the beluga whale, revealed alarming mismatches in demography and inhibited the detection of long runs of homozygosity^10^.

Despite the impact of reference bias on population genetic inference, few studies have examined how it may affect the characterization of demographic history and population recombination rates, which are increasingly popular targets of estimation. The gray fox (*Urocyon cinereoargenteus*) provides a compelling case study in this research. Gray foxes are widely distributed across North America, and despite their phenotypic similarity, genetic evidence suggests deep divergence between eastern and western lineages^11^. Previous gray fox genetic studies have relied on a domestic dog (*Canis lupus familiaris*) reference genome^12^, which poses challenges due to significant karyotypic differences between the two species^13,14^. In fact, despite most canid study’s reliance on the domestic dog reference genome, the clade contains a number of large karyotypic rearrangements, for example, dogs have 38 pairs of autosomal chromosomes, gray foxes have 32, and the Arctic fox (*Vulpes lagopus*) has 23-24.

The domestic dog, gray fox, and Arctic fox represent distinct evolutionary lineages within *Canidae*^15^. The gray fox represents the most basal lineage within the family, having diverged approximately 10 million years ago from all other living canids^15^. The remaining canids are comprised of three major clades: the red fox clade, the South American clade, and the wolf-like clade. The Arctic fox belongs to the red fox clade, which diverged from the wolf-like clade approximately 7-8 million years ago. The domestic dog is part of the wolf-like clade and diverged most recently, having split from other wolf-like canids about 3-4 million years ago.

There are also differences in the genome assembly sizes: the gray fox possesses the largest genome (2.66 Gb) compared to dog (2.48 Gb) and Arctic fox (2.35 Gb), some of which may be due to the sex of the individual selected for assembly. While GC content is relatively similar across all three species (∼42%), repeat content is variable, with the gray fox exhibiting the highest percentage (38.23%), followed by dog (33.67%) and Arctic fox (31.29%). These genomic differences are likely to bias downstream haplotype-based analyses that rely on synteny and linkage disequilibrium (e.g., demographic inference and recombination mapping). Despite this, most previous genomic research within *Canidae*—including studies on common (gray fox), near-threatened (Channel Island fox)^16,17^, and endangered (Ethiopian wolf)^18^ species—has been conducted by mapping to the domestic dog genome, primarily because it was the only canid reference genome available with annotations and had both high continuity and contiguity^15^.

The recently published gray fox reference genome^14^ provides us with a high-quality conspecific reference for alignment, variant calling, and ultimately for quantifying reference bias. Here, we re-analyze previously published WGS data from two populations of North American gray foxes^12^. We examine how inference of demographic histories and recombination landscapes differ when using the heterospecific domestic dog and Arctic fox genomes in contrast to the conspecific gray fox genome. To investigate the effects of reference bias on our analysis, we compare the underlying genetic variation, and the site frequency spectrum (SFS) estimated for each population using different reference genomes, as well as estimates of genetic diversity and differentiation. Finally, we examine the effect of reference bias on both coalescent and SFS based demographic inference, *F*_ST_ outlier scans for selection, as well as recombination rate inference.

## Results

We analyzed whole-genome resequencing data from 12 gray foxes, including six from the eastern U.S. and six from the western U.S. populations (Figure S1 and 1A). We examined reference bias by mapping these data to three reference genomes: the domestic dog genome CanFam4^19^, the Arctic fox genome^20^, and the conspecific gray fox genome^14^ (Figure 1B). Gray fox reads showed significantly higher mapping success to the conspecific reference genome compared to heterospecific genomes (99.7% vs. ∼99%; χ²_KW_=33.19, P<0.001, Table 1), with genes in unmapped regions enriched for sensory perception and immunity functions (details in supplementary text). Additionally, proper read pairing was higher with the conspecific reference (94.7%) vs. heterospecific references (89.4-90.3%), representing nearly 5% more properly paired reads (χ²_KW_=40.14, P<0.001).

**Figure 1.**
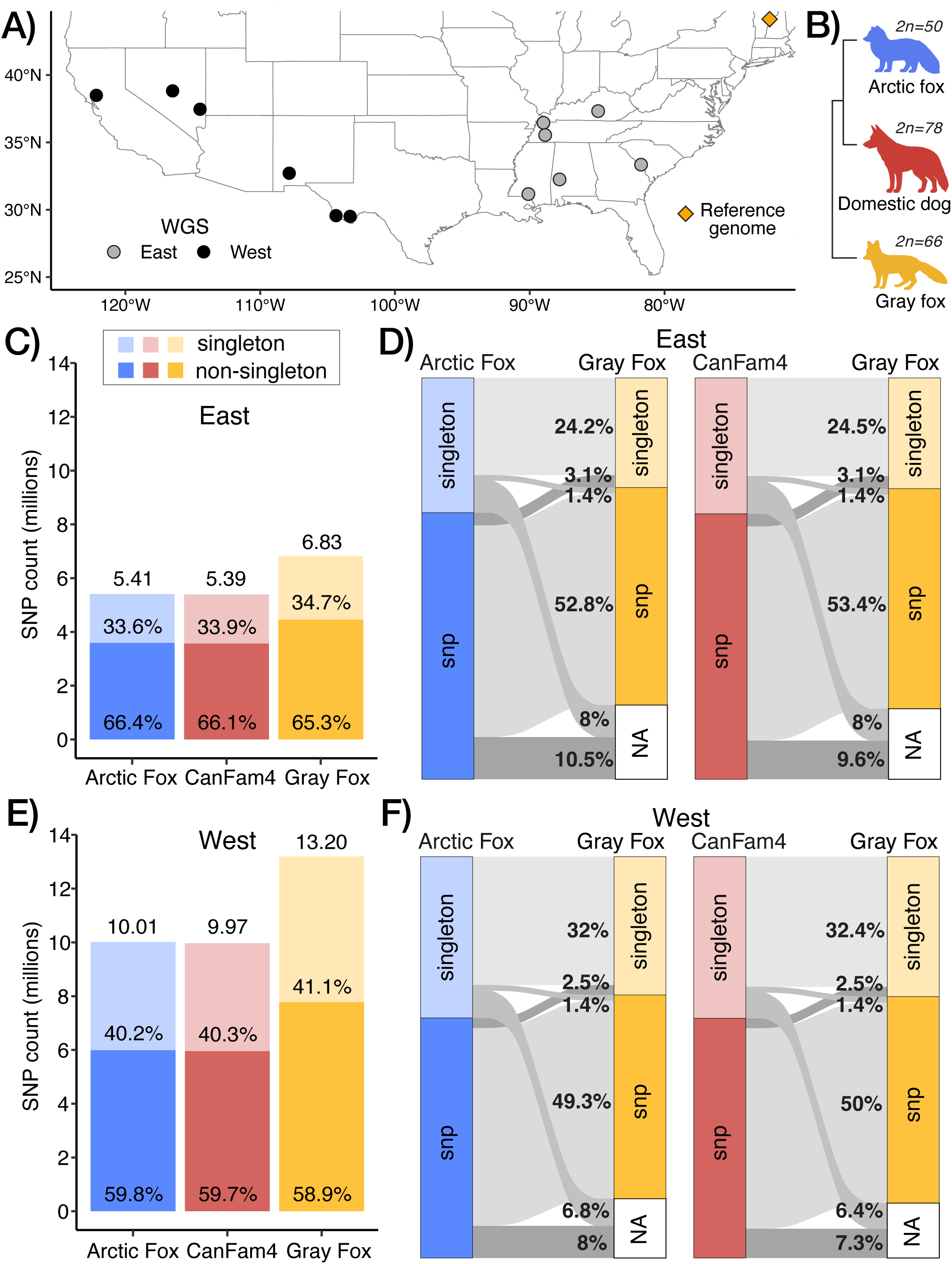
Reference bias influences variant detection. **A)** Sampling localities of gray fox genomes, with a star indicating the origin of the reference genome^14^ and circles representing WGS data^12^. **B)** Phylogenetic relationships of three canid reference genomes: Arctic fox (Vulpes lagopus, blue), domestic dog (Canis lupus familiaris, red), and gray fox (Urocyon cinereoargenteus, gold, most basal). Diploid chromosome numbers (2n) shown for each species. **C-F)** SNP counts and liftover classifications for eastern (C, D) and western (E, F) populations. Bar charts show total SNP counts (millions, above bars) divided into singleton (light) and non-singleton (dark) variants. Alluvial diagrams track variant reclassification when lifting over from heterospecific to gray fox reference genomes, showing transitions between singleton, SNP, invariant (“not snp”), and unmapped (“not map”) categories. Percentages show variant proportions and transition flows. Species-matched reference genomes yield higher effective population size estimates.

**Table 1.** The total number of successfully mapped reads, number of SNPs, and heterozygosity estimates (mean ± standard deviation) for each reference genome and population. The total number of reads (both mapped and unmapped) was 3,886,389,246 for the East and 1,043,628,176 for the West population.

| Genome | Population | Reads Mapped | Total SNPs | Heterozygosity (mean ± sd) |
| --- | --- | --- | --- | --- |
| Arctic fox | East | 3,836,962,124 | 5,407,943 | 0.181 ± 0.057 |
| CanFam4 | East | 3,848,140,971 | 5,392,973 | 0.181 ± 0.058 |
| Gray fox | East | 3,865,958,291 | 6,828,177 | 0.200 ± 0.051 |
| Arctic fox | West | 1032573693 | 10,012,122 | 0.163 ± 0.049 |
| CanFam4 | West | 1035433543 | 9,969,686 | 0.163 ± 0.049 |
| Gray fox | West | 1040317760 | 13,199,882 | 0.174 ± 0.047 |

**Table 2.**
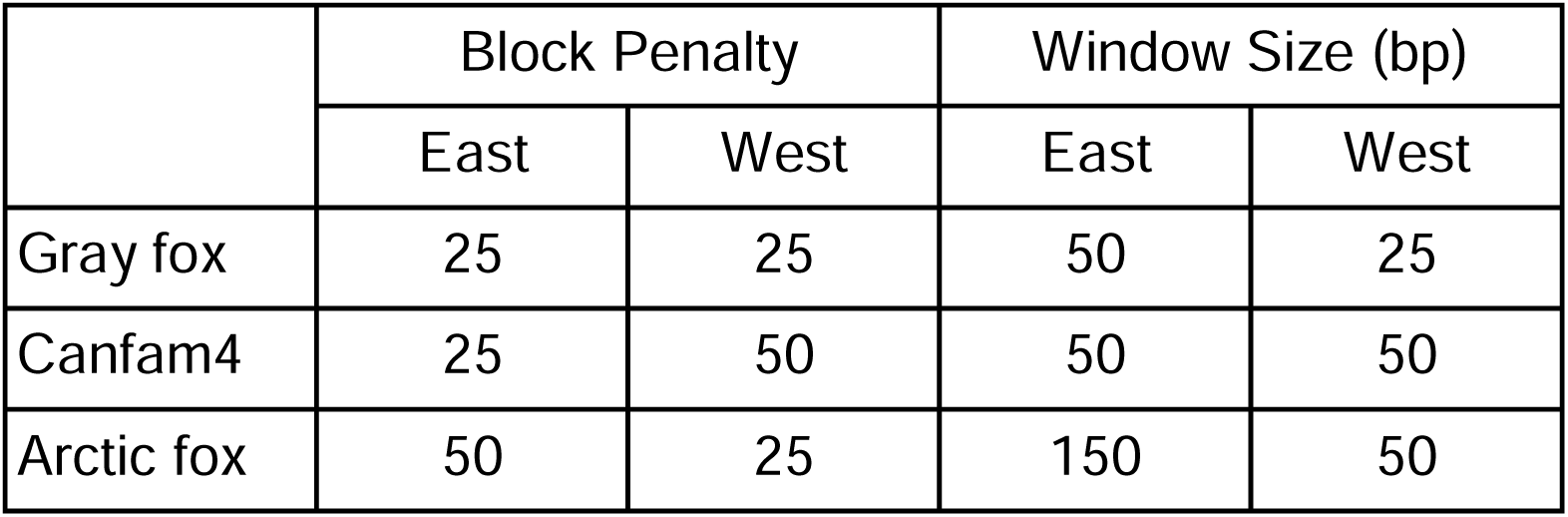
Pyrho hyperparameter settings for each reference genome and both populations.

|  | Block Penalty |  | Window Size (bp) |  |
| --- | --- | --- | --- | --- |
|  | East | West | East | West |
| Gray fox | 25 | 25 | 50 | 25 |
| Canfam4 | 25 | 50 | 50 | 50 |
| Arctic fox | 50 | 25 | 150 | 50 |

### A species-matched reference genome yields more SNPs and rare variants

We examined the impact of reference genome choice on the detection and characterization of genetic variation and allele frequency distributions in gray fox populations. Across all reference genomes, we detected more SNPs in the western population of gray foxes compared to the eastern population, with nearly double the number of SNPs identified in the west (Figure 1; Table 1). Notably, the gray fox reference genome consistently yielded more SNPs compared to the heterospecific references, with 26-32% more variants identified in each population (Table 1, Figure 1).

We identified more singletons (i.e., rare variants where an allele is only found once in the population sample) in the east and west using the gray fox reference (Figure 1,2). Specifically, the conspecific reference detected 33% more singletons (2.4M vs. 1.8M) in the eastern population and 35% more singletons (5.4M vs. 4M) in the western population compared to the heterospecific references. In addition, mean SNP depth varied slightly across genomes, with gray fox showing higher values (mean: east; 8.5, west; 8.7) compared to Arctic fox and dog (both east: 8.3, west: 8.6). Missing data on a per-site basis was lowest with the gray fox genome (east and west: 0.02), while the Arctic fox and dog genomes had higher missingness (east: 0.05, west: 0.03).

**Figure 2.**
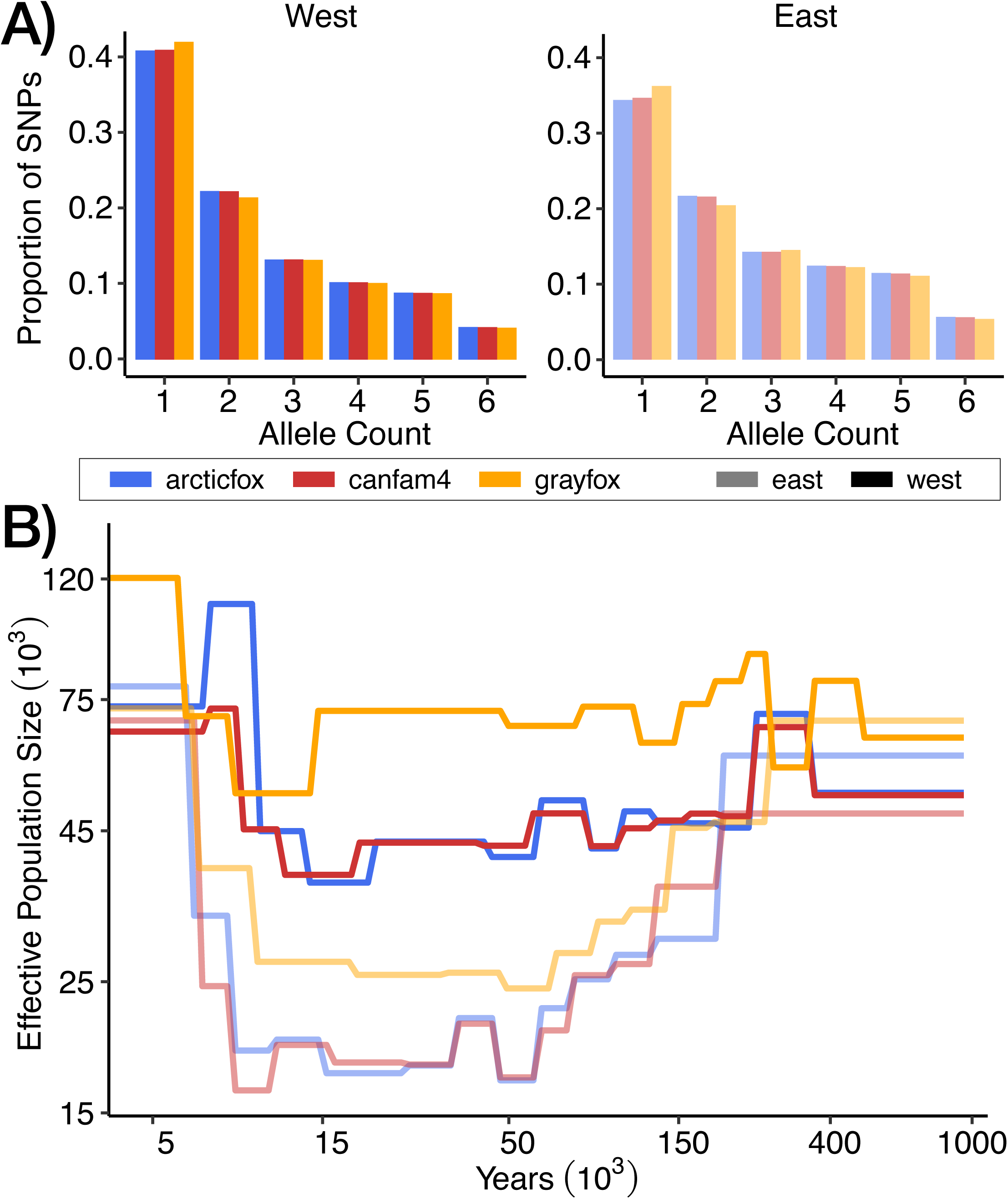
Reference bias influences the SFS and demographic trajectories. **A)** SFS for eastern and western populations show more singletons using the conspecific gray fox genome. **B)** Effective population sizes (y-axis) and years from present (x-axis) inferred with smc++ reveal discordant demographic histories of eastern (light) and western (dark) foxes resolved using the species-matched (gold) and heterospecific genomes (red and blue).

After lifting over SNPs in the heterospecific genomes to their corresponding positions in the gray fox genome, we found that approximately 80% of the variants identified using heterospecific reference genomes were also identified as variants when using the gray fox genome (Figure 1). Among these matching variants, non-singleton SNPs made up about 50-53%, while the remainder were singletons. The west population had a higher proportion of matching singletons compared to the east (∼32% vs. ∼24%), whereas the east population had a slightly higher proportion of matching non-singleton SNPs (∼53% vs. ∼50%). For the variants that did not lift over as SNPs, the majority (14-18%) were not identified as any type of variant in the gray fox genome. Of these, about half mapped to invariant sites, while the other half did not map at all (Table S1).

### A species-matched reference genome results in higher effective population size estimates

We inferred demographic histories using smc++^21^ and observed distinct population size trajectories when using different reference genomes for both the eastern and western populations. Across all genomes, the western population consistently showed higher effective population size (NL) estimates than the eastern population (Figure 2). The gray fox genome produced trajectories with smaller fluctuations in NL over time and generally higher NL estimates, particularly for the western population. In contrast, the Arctic fox and dog genomes revealed more variability, especially in the west, where the discrepancies between reference genomes were more pronounced. For instance, in the most recent time period, approximately 5,000-7,000 years ago, the gray fox genome indicated population growth in the west, although it should be noted that smc++ typically has higher uncertainty for estimates in the recent past, <6,000 years ago^21^. Meanwhile, the heterospecific genomes suggested a population decline, with NL estimates dropping below 75,000.

To evaluate whether removing variants that failed to map between species improves demographic inference accuracy, we compared smc++ trajectories before and after masking non-lifted-over SNPs in both heterospecific genomes and reciprocally in the gray fox genome. This approach made the demographic trajectories more similar across reference genomes (Figure S2). We also suspected that the total number of SNPs might affect demographic inference, so we down-sampled the gray fox SNPs to random subsets matching the number of variants available when using lifted-over variants from other genomes. These down-sampled trajectories were also similar to the lifted-over results, indicating that observed differences in demographic trajectories were driven not only by variant identity differences but also by the total number of informative variants available for analysis.

To evaluate how reference bias impacts demographic inference methods, we supplemented smc++ analysis (combining SFS with linkage disequilibrium in coalescent HMMs) with MSMC2 (coalescent HMM only) and Stairway Plot2 (SFS only) (Figure S3). Both MSMC2 and Stairway Plot 2 exhibited greater inconsistencies across reference genomes for the eastern population compared to the western population. MSMC2 inferred recent growth for the eastern population across all references, but heterospecific references exaggerated this growth, yielding eastern population sizes surpassing western ones—an outcome not observed with other methods and inconsistent with higher western diversity estimates. In contrast, Stairway Plot 2 revealed larger discrepancies in the eastern population during ancient times. For the western population, MSMC2 inferred recent growth only when using heterospecific references, whereas Stairway Plot 2 indicated stability across references, highlighting the varying sensitivities of demographic inference methods to reference bias. Despite variation across reference genomes, the methodological approach had a stronger influence on demographic inference outcomes. The disparities between methods highlight the critical importance of selecting appropriate demographic reconstruction methods, as method-specific biases can overshadow reference genome variation and underscores the need for careful method selection

### Recombination rates vary across reference genomes and populations

We detected significant differences in the distributions of recombination rates inferred using the different reference genomes in both populations (Anderson-Darling k-sample test, p<0.001; Figure 3). In the eastern population, recombination rates were lower when inferred using the Arctic fox genome compared to the gray fox genome. The average recombination rate was 0.448 cM/Mb with the Arctic fox genome, which is approximately 31% lower than the 0.650 cM/Mb observed with the gray fox genome. Conversely, when using the CanFam4 genome, the average increased to 0.872 cM/Mb, about 34% higher than the rate inferred with the gray fox genome. In the western population, recombination rates were higher when inferred using both heterospecific genomes compared to the conspecific gray fox genome. Specifically, the average recombination rate was 1.02 cM/Mb with the Arctic fox genome, approximately 13% higher than the 0.903 cM/Mb observed with the gray fox genome, and 0.989 cM/Mb with the CanFam4 genome, about 9.5% higher than the rate inferred with the gray fox genome. Overall, these differences were more pronounced in the east, where the heterospecific genomes led to both underestimation (Arctic fox) and overestimation (CanFam4) of recombination rates relative to the gray fox genome. In contrast, in the west, both heterospecific genomes consistently overestimated recombination rates compared to the gray fox genome, and the differences were less pronounced.

**Figure 3.**
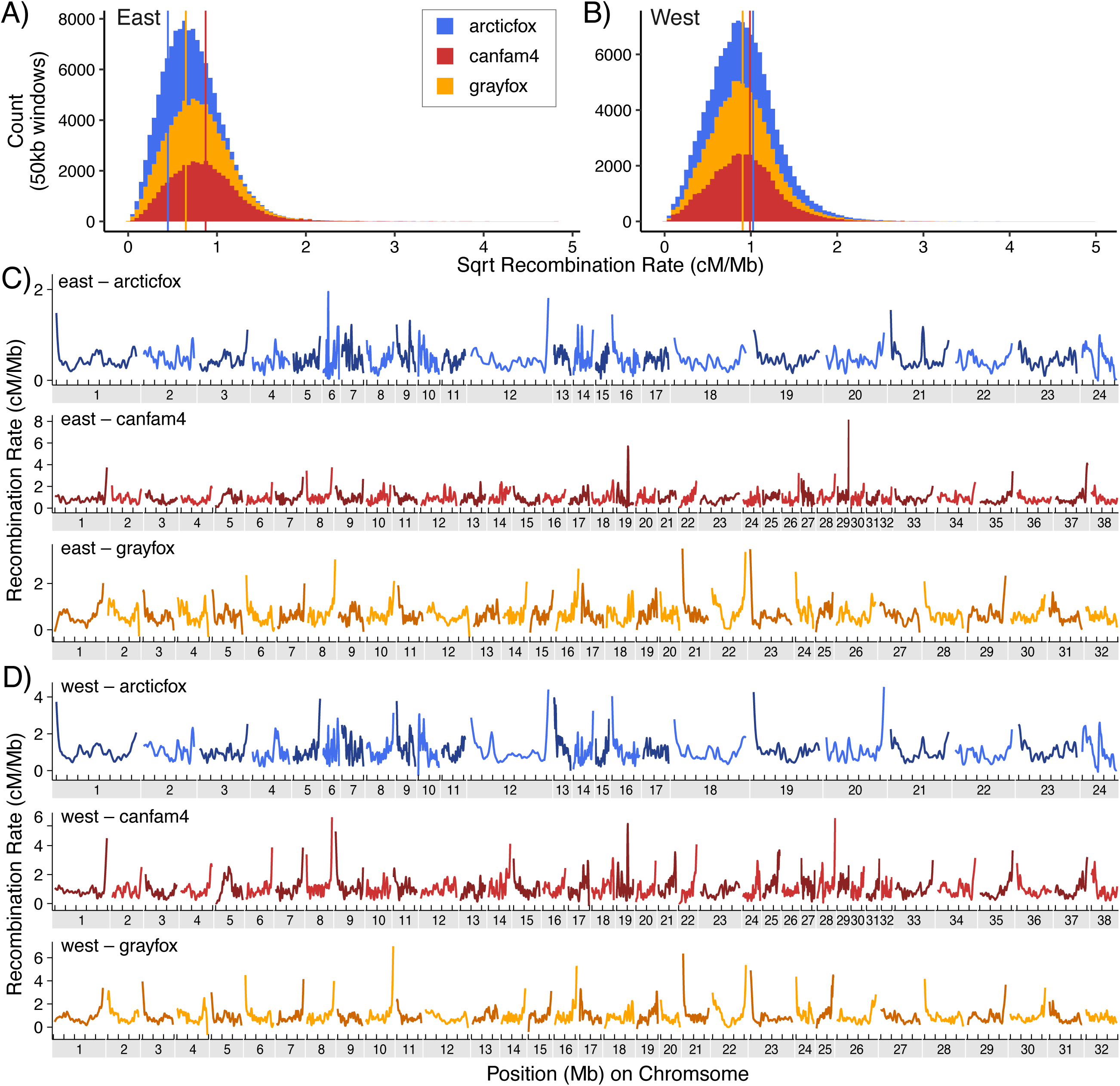
Recombination rate comparisons between reference genomes for gray fox populations. Histograms of the square root of recombination rates (cM/Mb) across 50 kb windows in (A) east and (B) west populations. Colors represent the three reference genomes used for recombination rate estimates, with vertical lines representing mean values. Loess-smoothed (span=0.1) recombination rates (cm/Mb) per chromosome computed over 50 kb windows in Eastern (C) and Western (D) gray foxes based on the three reference genomes.

Additionally, recombination landscapes based on each genome revealed substantial variation across 50 kb windows, with heterospecific reference genomes resulting in increased variability, particularly toward chromosome ends. In the eastern population, using the conspecific gray fox genome, recombination rates ranged from 0.000343 to 7.34 cM/Mb. However, when using the heterospecific Arctic fox genome, the maximum recombination rate doubled to 14.9 cM/Mb, and with the CanFam4 genome, it more than tripled to 23.4 cM/Mb. Similarly, in the western population, recombination rates estimated with the gray fox genome varied from 0.000598 to 11.6 cM/Mb. With the Arctic fox genome, the maximum recombination rate more than doubled to 24.9 cM/Mb, and with the CanFam4 genome, it increased to 14.5 cM/Mb. Compared to the gray fox genome, the heterospecific genomes resulted in higher maximum recombination rates, with a consistent trend of overestimation toward the ends of chromosomes, suggesting that heterospecific reference genomes can lead to an overestimation of recombination rates, especially at the higher end of the spectrum.

### Heterospecific genomes underestimate diversity and differentiation

We detected significant differences in estimates of average nucleotide diversity (π) in 50 kb windows in both gray fox populations based on the three reference genomes (Figure 4). In the eastern population, nucleotide diversity was lowest with the CanFam4 genome (π=0.000618), followed closely by Arctic fox (π=0.000625), while the gray fox genome yielded a higher estimate of π=0.000812 (χ²_KW_ = 6663.3, P<0.001). Similarly, in the western population, diversity was lowest using the heterospecific references (Arctic fox: π=0.00122, CanFam4: π=0.00121), while the gray fox genome produced a higher estimate of π=0.00164 (χ²_KW_=12409, P<0.001).

**Figure 4.**
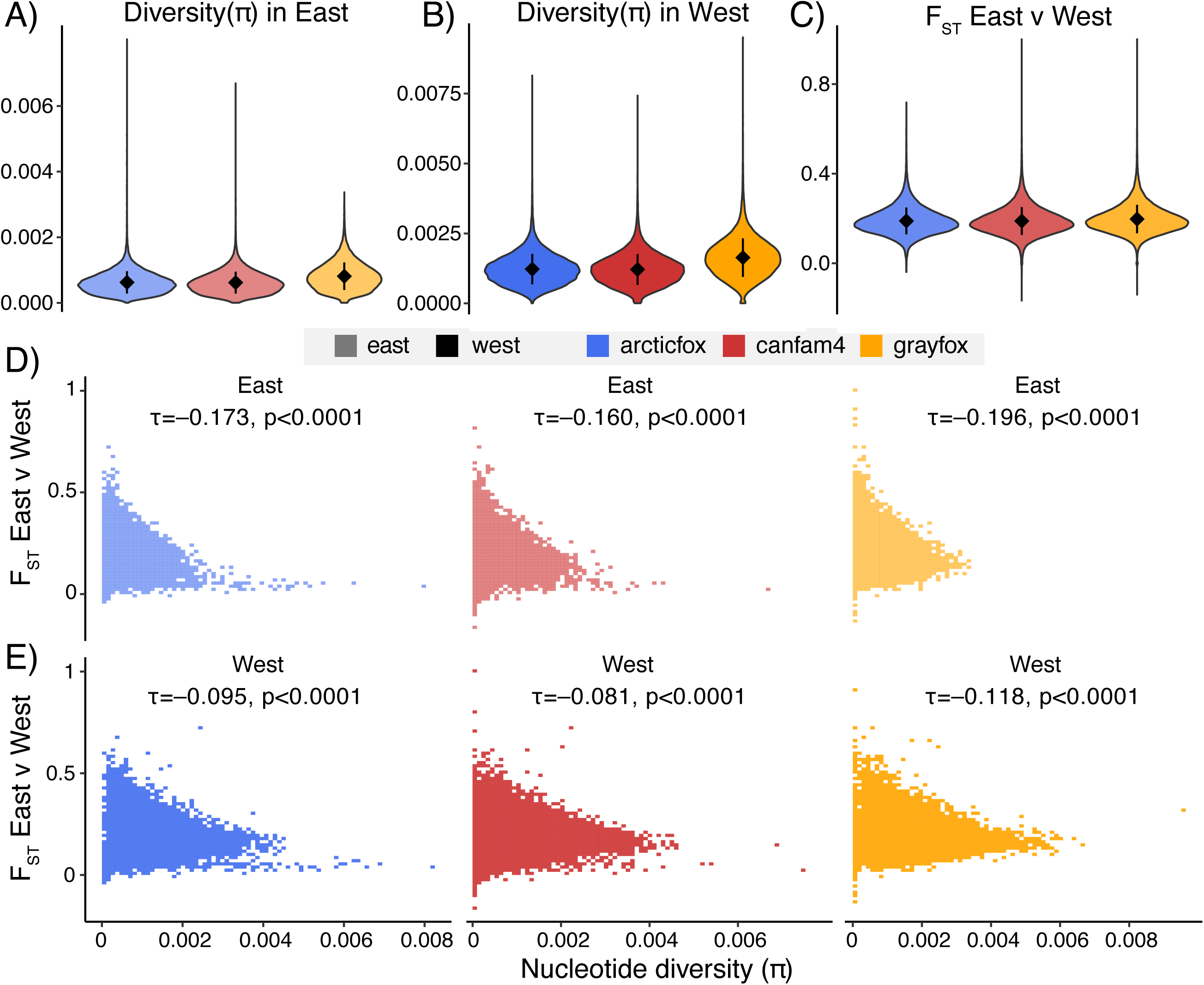
Estimates of diversity and differentiation are higher with gray fox reference in eastern and western gray fox populations. Nucleotide diversity (π) for eastern (A) and western (B) gray fox populations, and genetic differentiation (F_ST_) between the populations (C), in 50-kb windows based on three reference genomes. Violin plots with mean estimates and standard deviation error bars are shown. The correlation between diversity and differentiation across genomic windows for the east (D) and west (E) populations is illustrated. Kendall’s tau (τ) correlation coefficients are reported, showing significant negative correlations between π and F_ST_ in both populations across all reference genomes.

The variation in estimates of nucleotide diversity was slightly more evident in the west population, where π was approximately 1.34 times higher using the gray fox genome compared to the heterospecific references, while in the east, π was 1.31 times higher. While estimates of π were higher in the west than in the east across all reference genomes, the difference between populations was greater when using the gray fox genome, where diversity in the west was about 2.02 times higher than in the east. By contrast, diversity based on the Arctic fox genome showed a 1.95-fold difference between the west and east, and CanFam4 showed a similar 1.96-fold difference.

Estimates of genetic differentiation (*F*_ST_) between the eastern and western populations also varied depending on the reference genome. The heterospecific references produced identical *F*_ST_ values (mean *F*_ST_=0.189), while the gray fox genome resulted in a significantly higher mean *F*_ST_ of 0.197 (χ²_KW_=558.27, P<0.001). Additionally, the correlation between nucleotide diversity and *F*_ST_ was more strongly negative in the eastern population (τ=−0.173 to −0.196) compared to the western population (τ=−0.081 to −0.118), with the strongest correlations observed using the gray fox reference in both populations.

In both populations, mean Tajima’s D was lowest with the gray fox genome (−0.108 east, −0.356 west), compared to the Arctic fox (−0.056 east, −0.329 west) and CanFam4 (−0.063 east, −0.328 west) references (χ²_KW_ = 195.3 east, 144.4 west; P<0.001). Pairwise comparisons confirmed significantly lower Tajima’s D estimates with the gray fox genome (P<0.001), while estimates did not differ between the two heterospecific references (P=0.17 east, P=1.00 west). This indicates an excess of low-frequency polymorphisms when mapping to the conspecific genome, consistent with the higher number of singletons detected in the SFS based on the gray fox reference.

### Reference genome choice affects F_ST_ outlier detection

We defined outliers as windows with *F*_ST_ values exceeding three standard deviations above the mean for each reference genome, then matched these windows across reference genomes to identify shared and unique outlier regions. Across comparisons, we observed that heterospecific references consistently identified more than twice the number of unique outlier windows compared to the conspecific reference (Figure 5). Notably, the number of windows unique to each heterospecific reference was similar to the number of shared outliers between the heterospecific and conspecific references. In the Arctic fox and gray fox comparison, 148 shared outlier windows were identified, with 137 unique to the Arctic fox reference and 63 unique to the gray fox reference (Figure 5A). Between the Canfam4 and gray fox reference genomes, 165 outlier windows were shared, with 141 unique to Canfam4 and 61 unique to gray fox (Figure 5B). The majority of windows in both comparisons did not show elevated *F*_ST_ values in either reference (33,350 and 31,649 in the Canfam4 and the Arctic fox comparisons, respectively), reflecting our conservative method of outlier detection.

**Figure 5.**
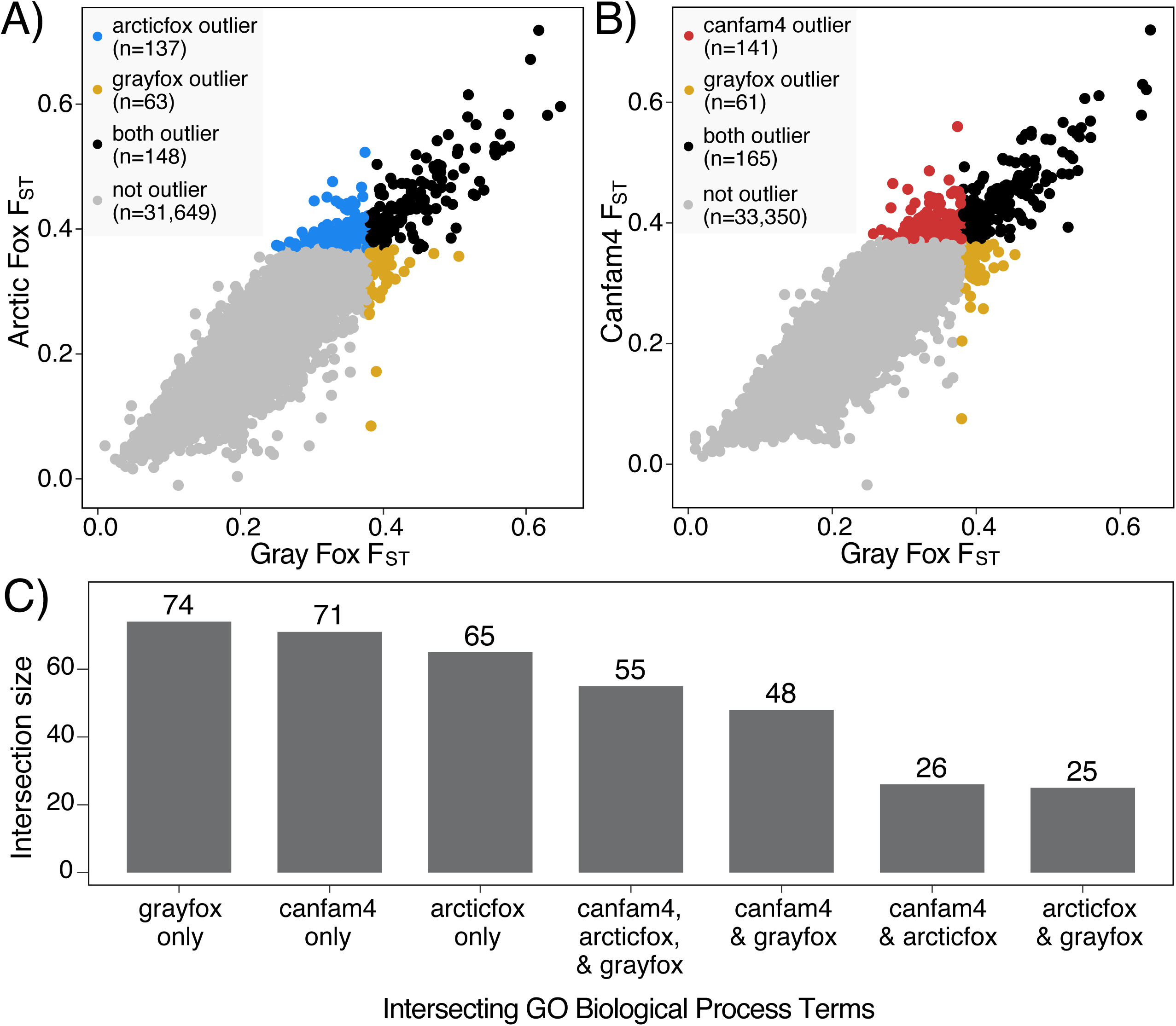
Reference-specific patterns revealed by F_ST_ outlier and functional enrichment analyses. (A) Scatter plot comparing F_ST_ values between gray fox and arctic fox references, with outlier windows unique to the arctic fox (blue) and gray fox (gold) highlighted. (B) Scatter plot comparing F_ST_ values between gray fox and Canfam4 references, highlighting outliers unique to Canfam4 (red) and gray fox (gold). For both (A) and (B), outliers shared between the two references are shown in black, while non-outlier windows are in gray. (C) UpSet plot of significant (p<0.05) GO biological process terms identified as enriched for each reference genome. Bars represent the intersection size (y-axis) of GO terms shared across reference genomes (x-axis).

Between the Arctic fox and Canfam4, we identified 191 shared outlier windows, 158 windows unique to the Arctic fox reference, and 42 windows unique to the Canfam4 reference, with 35,133 windows showing no elevated *F*_ST_ values in either reference (Figure S5). Thus, the Arctic fox and Canfam4 references shared slightly more outliers than either shared with gray fox (148-165), while the number of unique outliers for each heterospecific reference remained comparable to their gray fox comparisons.

### Functional enrichment patterns reveal reference-specific biological processes in F_ST_ outliers

The functional enrichment analysis of *F*_ST_ outliers revealed both distinct and overlapping gene ontology (GO) biological process terms across reference genomes (Figure 4C). Gray fox had the highest number of unique terms with 74, followed by Canfam4 with 71, and Arctic fox with 65, indicating that each reference captured distinct biological processes. Importantly, there were more unique terms than shared terms across references. Among shared terms, 55 were common to all three references. Notably, the overlap between gray fox and Canfam4 (48 shared terms) was larger than the overlap between Arctic fox and either gray fox (25 terms) or Canfam4 (26 terms), suggesting a closer alignment in identified biological processes between the gray fox and Canfam4 references.

Further, the unique terms identified by each reference genome suggested distinct thematic focuses (Figure S6). The Arctic fox reference highlighted processes related to cellular transport, secretion, and signaling, with terms like “positive regulation of catecholamine secretion” and “golgi to endosome transport,” indicating an emphasis on cellular signaling and developmental regulation. The Canfam4 reference captured growth, differentiation, and endocrine response, with terms such as “positive regulation of muscle cell differentiation” and “response to thyroid hormone,” pointing to processes involved in physiological development. In contrast, the gray fox reference emphasized nervous system function, immune modulation, and cell proliferation, reflected by terms like “positive regulation of nervous system process” and “calcineurin-mediated signaling.”

Gene Ontology (GO) enrichment analysis of genes located exclusively in Arctic fox reference-specific outlier windows revealed significant enrichment for terms related to muscle development and metabolism. The top biological process terms included “Muscle Organ Development” (GO:0007517), “Positive Regulation of Muscle Hypertrophy” (GO:0014742), and “Positive Regulation of Glycolytic Process” (GO:0045821). In contrast, genes in Canfam4 reference-specific outlier windows showed markedly different functional enrichment patterns, predominantly associated with chromosomal organization and cellular differentiation. The most significant terms included “Positive Regulation of Chromosome Organization” (GO:2001252), “Cell Differentiation in Spinal Cord” (GO:0021515), and “Regulation of Chromosome Separation” (GO:1905818). These distinct functional enrichment patterns demonstrate how reference genome choice can lead to substantially different biological interpretations. The Arctic fox reference-specific outliers suggest selection on genes involved in muscle development and energy metabolism, which could be interpreted as adaptations related to locomotion, hunting behavior, or thermal regulation. Conversely, the Canfam4 reference-specific outliers point toward selection on chromosomal organization and cellular differentiation pathways, potentially indicating differences in developmental processes or cell cycle regulation.

## Discussion

Reference genomes are essential tools in genetic studies, serving as coordinate systems for annotating sequence features and comparing individuals and populations. However, many non-model organisms lack annotated chromosome-level assemblies^22^, forcing researchers to use genomes from closely related species^23,24^, which can compound reference bias. Reference bias has been previously studied across the tree of life^5,6,10,25–28^ both within species^26,27^ and between species^6^. In this study, we demonstrate that mapping to a heterospecific reference genome significantly impacts population genomic inferences. This leads to discrepancies in effective population size estimates, recombination rates, measures of genetic diversity, differentiation, and selection signals. Our findings highlight the importance of using conspecific genomes for accurate evolutionary inferences.

Although the average sequencing depth showed only minor discrepancies using a discordant reference, we observed significant differences in both the percentage of reads mapped and the proportion of properly paired reads. Reads are not properly paired when the read orientation is incorrect, one read does not map at all, and/or the gap between the reads are of an unexpected size. Although thresholds for gap size and mismatch rates can be changed during read alignment, most studies use inconsistent filters at this stage of the pipeline, including retaining only reads with a certain MAPQ score, retaining proper pairs, or only retaining the primary alignment^29^. Likely, the number of properly paired reads decreases when the gray fox is mapped to the heterospecific genomes because of structural variation between the genomes, some of which may be smaller indels, and some of which may reflect the large chromosomal rearrangements between the species. Following common practice, we retained improperly paired reads, which may explain downstream variant detection effects, though selective filtering could potentially mitigate this.

To exemplify the inconsistency of pipelines and filters applied, we screened five papers which used the domestic dog as a reference genome for analyses of a heterospecific canid taxa and found that four papers did not remove improperly paired reads before variant calling^12,16,18,30^, while one paper did remove improperly paired reads^17^, and their remaining mapping and filtering choices had almost no overlap. Despite many of these studies filtering on MAPQ, mapping quality is assigned for an individual read by BWA^31^, and not as a pair (i.e., the MAPQ is ambiguous to the read being properly paired). GATK automatically filters reads with a MAPQ of 10 or less during variant calling and other filters such as genotype quality (GQ; accuracy of the particular variant call at that site) and QUAL (how confident the variant caller is that there is variation at that site) are also likely to affect the resulting variant calls. Though we do not explore these as part of this work, a systematic investigation of these filters and their downstream impacts on mapping and variant calling, especially in the case of divergent references, is highly needed.

We found that the conspecific gray fox genome provided a more accurate representation of genetic variation, revealing more SNPs and a more reliable SFS compared to the heterospecific genomes. Heterospecific references underestimated the presence of low-frequency alleles, which can have profound implications for downstream analyses. For example, underestimating rare variants can lead to failure to detect recent population expansion in demographic inference, as the characteristic excess of rare alleles that signals expansion would be artificially reduced. Additionally, an underestimation of genetic load can occur when assessing deleterious mutations, since many harmful variants tend to be maintained at low frequencies due to purifying selection. Furthermore, recombination rate estimates in population-scale maps can become distorted due to missing variants, as these estimates rely on accurate measurement of linkage disequilibrium patterns between neighboring loci.

The higher mean SNP depth and lower missing data rates with the gray fox reference further support its superior performance in variant detection, indicating a more accurate and comprehensive alignment that reduces the likelihood of false negatives. Approximately 20% of the variants identified with heterospecific references were incorrectly identified when lifted over to the gray fox genome (Figure 1). Of these, ∼5% were misclassified (singleton vs. non-singleton), and ∼15% were unrecognized as variants, with half mapping to monomorphic sites and half failing to map (Table S1). The 20% discrepancy likely results from sequence divergence between species, leading to alignment errors or misinterpretation of homologous regions. The higher proportion of matching non-singleton SNPs suggests that common variants are more consistently detected across references, while rare variants are more likely to be missed when using heterospecific genomes. When estimating nucleotide diversity (π), the gray fox reference produced higher values in both populations, indicating that heterospecific genomes can underestimate genetic diversity within populations. The greater difference in π between the western and eastern populations observed with the gray fox genome also suggests that the conspecific reference provides a more sensitive measure of genetic diversity.

Having performed the first demographic inference for the gray fox with a conspecific reference genome, we sought to compare the eastern and western gray fox populations to gain deeper insights into their population histories. These two populations had quite distinct demographic trajectories, with the western population showing overall larger and stable population size with no recent growth. In contrast, the eastern population showed a stable population with some recent growth. Genetic diversity was significantly higher in the western gray fox population compared to the eastern population, suggesting differing evolutionary pressures and histories in these lineages. Our results differ from previous studies^12^ of these populations, which mapped to CanFam3.1 and used PSMC on a single individual to infer the ancient demographic histories of the populations. Specifically, we infer smaller and more stable populations throughout time, with less of a disparity between east and west population sizes. Both our study and the previous study identified the same pattern of the western population being more diverse than the eastern population for most timepoints, though the magnitude of our diversity estimates is much lower than previous estimates^12^. We also did not detect a recent decline in the west, which was found only when mapped to heterospecific references in our analysis as well as in previous work^12^.

We used multiple methods for demographic inference to determine if coalescent or SFS-based approaches were more robust to reference bias. All methods showed that heterospecific genomes consistently underestimated effective population sizes compared to the conspecific reference. The conspecific reference produced estimates that align more closely with expectations based on species biology and similar studies in other mammals^32^. Importantly, heterospecific genomes produced demographic trajectories inconsistent with the conspecific reference throughout most of the inferred history, regardless of population or inference method (Figure S3). This inconsistency is particularly worrying in the case of species threatened with extinction, because approximately 99% of these species do not have a reference genome^33^ and researchers will ultimately be forced to map to a heterospecific reference. When researchers must use heterospecific references, several strategies can mitigate reference bias. For instance, using end-to-end alignment methods that require the entire read to align reduces bias at indels, compared to local aligners that allow parts of reads that do not align to be ignored (i.e., soft clipped)^34^. Using local realignment methods that ensure consistent gap placements across all reads covering the same region, and comparing results across different filtering thresholds can also help reduce bias^34^. Additionally, constructing consensus references or using multiple population reference genomes in a “reference flow” approach effectively reduces bias while requiring fewer computational resources than graph-based methods^35^.

Since pedigree-based recombination estimates are difficult to obtain and have low resolution, researchers typically use population-based methods like pyrho^3^ to infer recombination maps that account for demographic histories. Focusing on the recombination rate estimates inferred with the species-matched reference, we found that landscapes of recombination differed notably between the eastern and western populations, with the west consistently exhibiting higher recombination rates, particularly at chromosome ends. A recombination landscape that is fairly stable with peaks towards the end of the chromosome has been previously observed in dogs^36^ and is particularly interesting given that the family *Canidae* has a psuedogenized copy of positive-regulatory domain zinc finger protein 9 (PRDM9)^14,37^. PRDM9 is known to initiate meiotic recombination by specifying the locations of double-strand breaks and creating recombination hotspots. Previous work highlighted how its pseudogenization in *Canidae* contributes to observed recombination patterns in dogs, which can be used to identify the directionality of chromosomes. Chromosome directionality has yet to be established for the new reference genome, so future work could take advantage of the result to identify directionality of chromosomes in the gray fox.

Conversely in the case of heterospecific reference genomes, where demographic trajectories were quite discordant from the conspecific reference, the inferred recombination maps and rates differed significantly in both populations. In the eastern population, recombination rates were underestimated with the Arctic fox genome and overestimated with the dog genome compared to the gray fox genome, while in the west, both heterospecific genomes overestimated recombination rates, with less pronounced differences. The inconsistent direction of bias— where the Arctic fox genome underestimates recombination rates in the eastern population but overestimates them in the western population compared to the gray fox genome—suggests that population-specific patterns may influence how reference bias manifests. This variability can complicate mitigation efforts, as the potential sources of bias differ between populations, making uniform corrections ineffective and highlighting the complexities in addressing reference bias when it varies between populations. In addition, recombination landscapes based on heterospecific genomes showed greater variability, particularly toward chromosome ends, with higher maximum rates than those inferred using the gray fox genome. These patterns highlight how reference genome choice can introduce substantial bias, affecting both the magnitude and distribution of inferred recombination rates.

Potential sources of bias for the inferred recombination rate include chromosomal synteny and spurious singletons. In the context of synteny, previous work has shown that there have been large scale chromosomal rearrangements within the evolutionary history of these species^14^. These karyotype differences have resulted in less than half of the dog and Arctic fox chromosomes being syntenic with gray fox chromosomes (Figure S8; Table S2). Thus, when a recombination map is estimated using a heterospecific genome with karyotypic differences as a reference, we are ultimately disrupting linkage patterns and inferring false recombination events across multiple chromosomes. To investigate this, we examined whether differences in recombination rates between each heterospecific reference and the conspecific reference varied between syntenic and non-syntenic chromosomes but found no difference in the mean recombination rates in either population or genome comparison (Figure S9). The observed inflation in recombination rates could result from disrupted linkage patterns caused by lack of synteny, increased sequence divergence leading to spurious singletons, or an interaction of these factors. Disentangling these contributions is challenging because they are inherently interconnected: synteny disruptions and sequence divergence both affect how variants are identified and mapped, while spurious singletons may arise as an artifact of these processes, obscuring their individual effects. In syntenic regions, pyrho^3^ would infer inflated recombination rates when there are spurious singletons and other incorrectly mapped variants being interpreted as evidence of recombination events. The misclassification and misidentification of ∼20% of the variants as a result of reference bias (Figure 1) is likely what led to skewed estimates of recombination rates, especially in regions of the genome where alignment is poor. The observation of non-syntenic regions and lower reference genome quality introducing bias was also captured in recent work examining an improved olive baboon reference genome^38^. Finally, we examined estimates of *F*_ST_ and used the genome-wide distribution of *F*_ST_ to perform an outlier-scan to detect genes that are potentially under selection. We found that estimates of *F*_ST_ were higher when using the gray fox genome compared to the values obtained from heterospecific references. We also observed a stronger negative correlation between π and *F*_ST_ when using the gray fox reference, suggesting that regions of low diversity correspond to areas of high differentiation. The discrepancy among overall π and *F*_ST_ values across reference genomes is not concerning in terms of magnitude, suggesting that genome-wide averages are likely sufficient for broadly characterizing diversity within, and differentiation between, populations. However, studies typically go beyond genome-wide averages, examining variation in *F*_ST_ across the genome to identify regions of high differentiation, which may indicate the presence of genes under selection or regions contributing to reproductive isolation.

Importantly, we found that the choice of reference genome substantially affected *F*_ST_ outlier detection. Heterospecific references identified more than twice the number of unique outlier windows compared to the conspecific reference. This discrepancy could be due to alignment issues, where regions not well conserved between species may erroneously appear as highly differentiated regions. The limited overlap of outlier windows between references raises concerns about the reliability of using heterospecific genomes for detecting genomic regions under selection. The distinct GO terms associated with each reference further illustrate how reference genome choice shapes the inferred biological processes underlying *F*_ST_ outliers. For instance, unique GO terms in the Arctic fox reference emphasized cellular signaling and developmental regulation, while Canfam4 highlighted growth and endocrine response, and the gray fox reference captured nervous system function and immune modulation. These differences underscore the biological biases introduced by reference genome choice. In sum, using a heterospecific reference genome would inflate false positives for both the *F*_ST_ outliers, genes, and biological pathways that may be under selection. Thus, using a conspecific reference would certainly provide more accurate insights into the biological processes influencing genetic differentiation within and between species.

This study highlights the critical impact of reference genome choice on population genomic inferences and emphasizes the value of conspecific genomes in uncovering accurate evolutionary histories. Quantifying reference bias using *Canidae* has implications for some of the world’s most endangered species, such as the Ethiopian wolf (*Canis simensis*) and the African wild dog (*Lycaon pictus*), and more common species, like the gray fox and coyote (*Canis latrans*). Reference bias may also affect our understanding of historic hybridization and introgression, a phenomenon common in the canid clade^30,39^. Furthermore, genomic data are increasingly being used to inform conservation management plans. Measures such as adaptive capacity^40^ and differential adaptations between populations^41^ are being used to make recommendations regarding translocations and rewilding. Our selection scans suggest that these inferences are skewed by reference bias when divergent reference genomes are used.

Our results underscore the necessity of using conspecific reference genomes in conservation genetics and evolutionary studies, particularly for accurately understanding divergence and diversity in non-model organisms. However, we acknowledge that developing high-quality reference genomes for all species is not always feasible due to resource constraints. In such cases, careful consideration should be given to the phylogenetic proximity and degree of synteny of the available reference genomes. Our work demonstrates that even closely related heterospecific references may not adequately capture the genetic landscape of the target species. Future studies should focus on improving reference genome assemblies for non-model organisms, leveraging advancements in sequencing technologies and assembly algorithms. Additionally, methodologies that are less reference-dependent, such as reference-free variant calling or pangenomic approaches, may help mitigate some of the biases introduced by heterospecific references.

### Limitations of the Study

Our work provides important insights into reference bias but has limitations. First, the gray fox reference genome comes from an east coast individual, potentially introducing bias that future work could mitigate computationally or by generating a west coast reference. Second, we quantified reference bias in *Canidae*, a mammalian clade with notable karyotype shifts. Despite these shifts, *Canidae* species are relatively young, likely moderating reference bias compared to more divergent taxa. More divergent non-mammalian taxa are likely to experience stronger reference bias. Since non-mammalian species comprise most of the 99% of species without reference genomes, this highlights the broader importance of addressing reference bias.

However, karyotype differences occur even between closely related species, making *Canidae* useful for studying the interplay between karyotype shifts and reference bias. Additionally, our mapping and filtering parameters, while representing common practices, may influence the magnitude of reference bias observed. Further analysis is needed to examine how these choices interact with reference bias and impact other genomic analyses, including inference of introgression patterns and linkage disequilibrium, which were beyond the scope of this study. This study does not address reference bias in ancient DNA. Previous research has shown that, in addition to DNA quality, reference bias is influenced by factors such as sequence length, genetic divergence^43,44^, the alignment tool used^43^, and map quality filtering^43,45^. Genetic summary statistics require careful interpretation in ancient DNA studies, where both human and non-human extinct lineages will be mapped to divergent interspecific references. The additional temporal component of ancient DNA will likely amplify the biases observed here.

### Conclusion

We demonstrate that using a conspecific reference genome enhances the inference of population size histories and recombination rates, improves the detection of genetic variation, provides more accurate estimates of nucleotide diversity and genetic differentiation, and influences the biological interpretation of genomic data. These findings highlight the broader implications for genomic research in other non-model organisms and stress the need for continued efforts in generating high-quality, species-specific genomic resources. Considering these biases, our work demonstrates that results generated from mapping to a highly divergent reference genome should be interpreted with caution.

## Acknowledgements

We want to thank Doc Edge, Joshua Schraiber, and Jeffrey Spence for insightful and helpful conversations during this work. J.A.M was supported by the startup funds from Dornsife College of Letters, Arts and Sciences through the Department of Quantitative and Computational Biology and the USC WiSE Gabilan Assistant Professorship. M.G. was supported by USC Provost’s Fellowship.

## Data accessibility and benefit-sharing

GitHub code for filtering can be found here https://github.com/ellieearmstrong/Gray_Fox_2023/tree/main/filtering and analyses can be found at https://github.com/makopyan/fox/.

## Author contributions

M.A. and J.A.M. conceived of the project. J.A.M., M.A., and M.G. performed data processing and analyses. E.A. conducted mapping and generated variant call files used for analyses. J.A.M., M.A., E.A., and M.G. wrote the manuscript. All authors approved of the final manuscript.

## STAR Methods

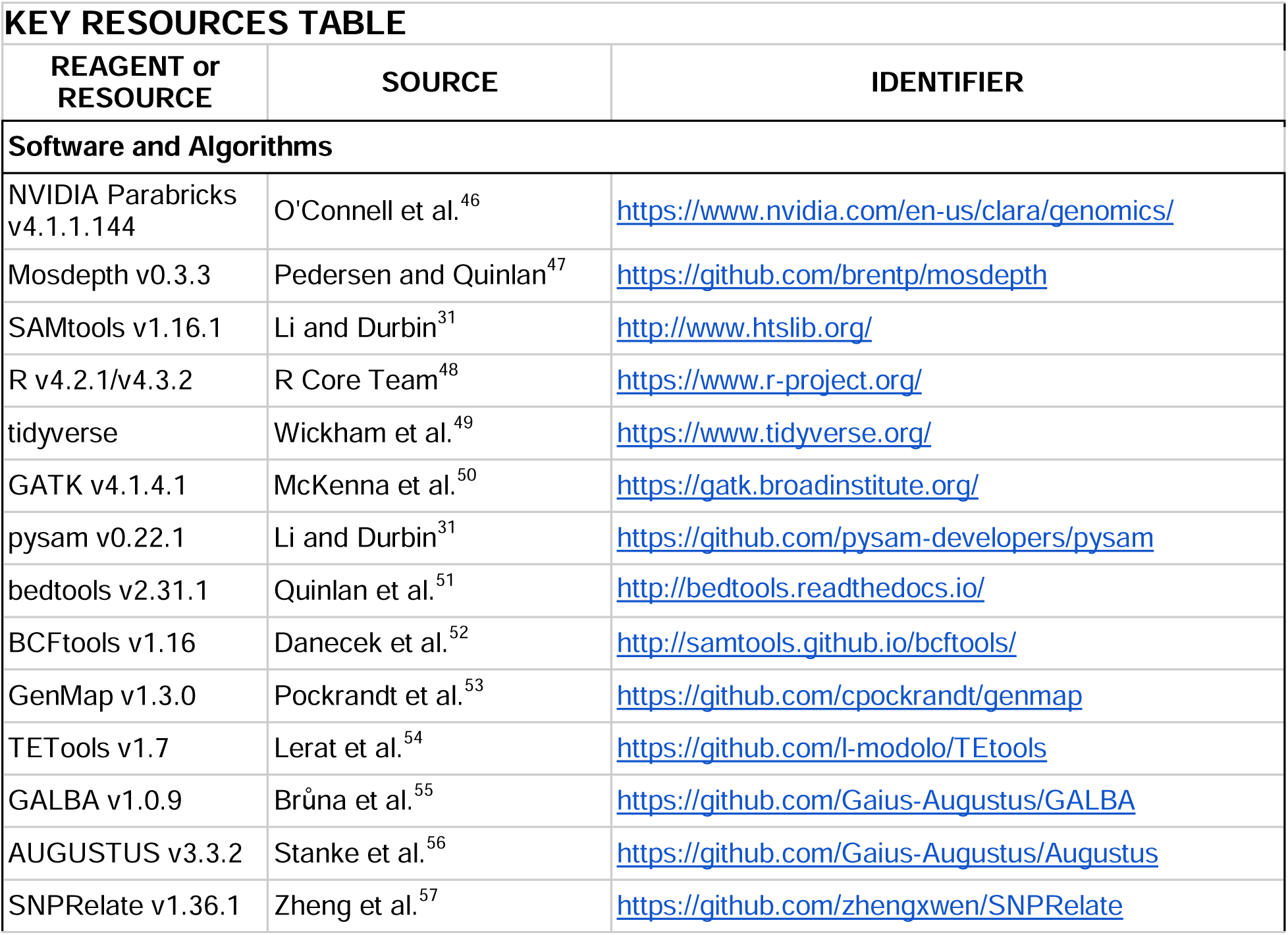
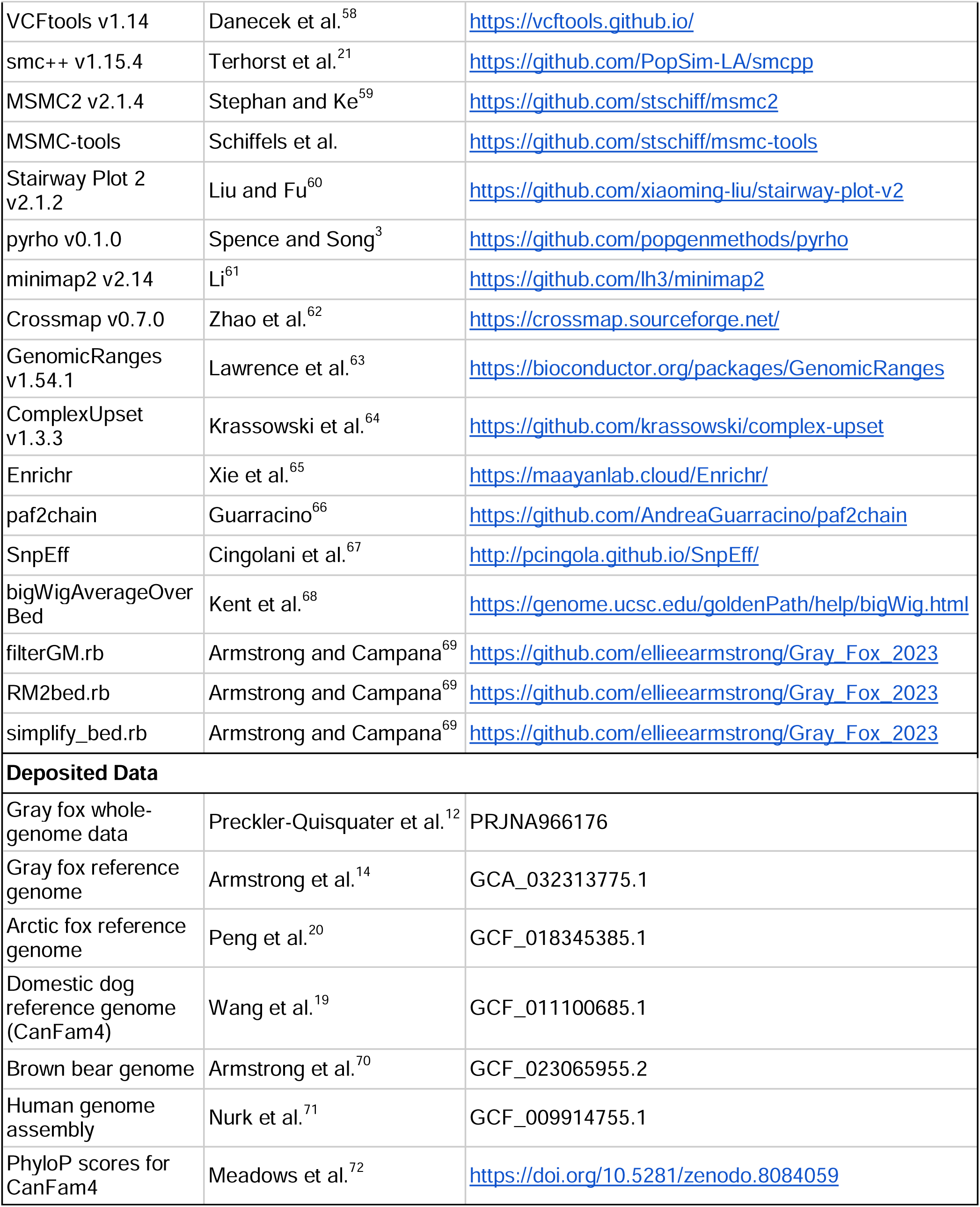

