## SupplementaryFigsAndText for "Reference genome choice compromises population genetic analyses"

### Supplementary Methods

#### Whole genome sequence data and reference genomes

To investigate the impact of reference bias on population genomic inference of diversity, demography, selection, and recombination in gray foxes, we obtained whole-genome resequencing data for 41 gray foxes sampled across North America (Figure S1) from Preckler-Quisquater et al.[^1^](https://app.readcube.com/library/bc53ac12-924a-4977-b177-875ccbf08ca1/all?uuid=9240579200031448&item_ids=bc53ac12-924a-4977-b177-875ccbf08ca1:9d9c5b74-25b4-4d11-b82a-6467769efd3f), in which the data were mapped to the domestic dog reference genome CanFam3.1[^2^](https://app.readcube.com/library/bc53ac12-924a-4977-b177-875ccbf08ca1/all?uuid=772364581295638&item_ids=bc53ac12-924a-4977-b177-875ccbf08ca1:9b985107-498e-415e-a4f3-ad276067a49e). We re-mapped the data to three reference genomes: (1) Arctic fox[^3^](https://app.readcube.com/library/bc53ac12-924a-4977-b177-875ccbf08ca1/all?uuid=20619428479684&item_ids=bc53ac12-924a-4977-b177-875ccbf08ca1:d6c8cdc2-b55c-4910-9cde-bc2b0d508642), (2) a more contiguous updated version of the dog genome CanFam4[^4^](https://app.readcube.com/library/bc53ac12-924a-4977-b177-875ccbf08ca1/all?uuid=25445926241542094&item_ids=bc53ac12-924a-4977-b177-875ccbf08ca1:a10e08eb-cd9b-4502-ae02-d5757a7bfff3), and (3) our recently published gray fox reference genome[^5^](https://app.readcube.com/library/bc53ac12-924a-4977-b177-875ccbf08ca1/all?uuid=8306136333047993&item_ids=bc53ac12-924a-4977-b177-875ccbf08ca1:5ddd40e9-72a1-497b-8efe-aec57a4ebcd8), based on an individual sampled in Vermont (Figure 1A). We identified syntenic and non-syntenic chromosomes between species using information from Graphodatsky et al.[^6^](https://app.readcube.com/library/bc53ac12-924a-4977-b177-875ccbf08ca1/all?uuid=14336281505805515&item_ids=bc53ac12-924a-4977-b177-875ccbf08ca1:3b7511b5-ccf8-48be-ab93-0d4589f76e2e) and through alignments generated in Armstrong et al.[^5^](https://app.readcube.com/library/bc53ac12-924a-4977-b177-875ccbf08ca1/all?uuid=19946136372573842&item_ids=bc53ac12-924a-4977-b177-875ccbf08ca1:5ddd40e9-72a1-497b-8efe-aec57a4ebcd8). All putatively syntenic chromosomes from Graphodatsky et al.[^6^](https://app.readcube.com/library/bc53ac12-924a-4977-b177-875ccbf08ca1/all?uuid=7033347805717938&item_ids=bc53ac12-924a-4977-b177-875ccbf08ca1:3b7511b5-ccf8-48be-ab93-0d4589f76e2e) were then manually interrogated using the genomic alignments from Armstrong et al.[^5^](https://app.readcube.com/library/bc53ac12-924a-4977-b177-875ccbf08ca1/all?uuid=3279731028227424&item_ids=bc53ac12-924a-4977-b177-875ccbf08ca1:5ddd40e9-72a1-497b-8efe-aec57a4ebcd8). Any putatively syntenic chromosome that mapped to more than one chromosome in either species was removed and considered non-syntenic to be as conservative as possible.

#### Mapping and variant calling

We mapped whole-genome data from gray fox individuals to each of three reference genomes (see above) using identical pipelines. Sequence data was mapped using NVIDIA Parabricks v4.1.1.1[^7^](https://app.readcube.com/library/bc53ac12-924a-4977-b177-875ccbf08ca1/all?uuid=5552228818184398&item_ids=bc53ac12-924a-4977-b177-875ccbf08ca1:5ad8874f-5594-473c-9b6a-584053eb1cad) *fq2bam* with the provided reference and default parameters. We estimated depth and other mapping statistics using Mosdepth v0.3.3[^8^](https://app.readcube.com/library/bc53ac12-924a-4977-b177-875ccbf08ca1/all?uuid=5407485857783736&item_ids=bc53ac12-924a-4977-b177-875ccbf08ca1:e6588e1a-c1b7-4e2c-b059-5ea1543cd00b) and SAMtools v1.16.1 *flagstat*, respectively. We tested for differences in these measures using pairwise.wilcox.test with p.adjust.method = “none” in R v4.2.1[^9^](https://app.readcube.com/library/bc53ac12-924a-4977-b177-875ccbf08ca1/all?uuid=5860347283591886&item_ids=bc53ac12-924a-4977-b177-875ccbf08ca1:6d003005-8cc7-4751-b838-df235f4833d9). Resulting BAM files were individually run through Parabricks *HaplotypeCaller* with the ‘--gvcf’ flag in order to emit both variant and invariant site calls. Subsequently, GATK v4.1.4.1[^10^](https://app.readcube.com/library/bc53ac12-924a-4977-b177-875ccbf08ca1/all?uuid=9396735198988391&item_ids=bc53ac12-924a-4977-b177-875ccbf08ca1:0ff45392-8020-453c-a84e-ea1953249b86) *GenomicsDBImport* was used to import the single sample VCFs prior to joint genotyping with the ‘-all-sites’ flag to retain both variant and invariant sites. Autosomal chromosomes were provided as intervals, excluding sex chromosomes and unlinked scaffolds for downstream analyses. Finally, we ran GATK *GenotypeGVCFs* with the ‘--all-sites’ flag to produce final gVCF files. Files were combined using BCFtools v1.16[^11^](https://app.readcube.com/library/bc53ac12-924a-4977-b177-875ccbf08ca1/all?uuid=10358358458507222&item_ids=bc53ac12-924a-4977-b177-875ccbf08ca1:9c355c42-03ca-4b22-89f6-9a90d97aad44) *concat* to create genome-wide variant calls. Read mapping counts were generated using pysam v0.22.1[^12^](https://app.readcube.com/library/bc53ac12-924a-4977-b177-875ccbf08ca1/all?uuid=8999478975902144&item_ids=bc53ac12-924a-4977-b177-875ccbf08ca1:b25637a1-1f2c-44f5-a2b2-2d963c4738ce) and custom scripts.

#### Variant filtering

Both repetitive and low-mappability regions were filtered using the BCFtools *view* command and providing the described bed file after the ‘-T ^’ flags. We then assessed quality statistics using BCFtools *query* with the flags ‘-f’ and pulled the statistics for allelic number (AN) and depth (DP). We calculated the mean and interquartile range for allelic number and depth for these statistics for each genome. Subsequently, we used the BCFtools *filter* command with the ‘-i’ flag to include variants which had more than 90% of the maximum AN value (2* number of samples), a quality score greater than 30 (‘QUAL >= 30’), and a depth that was greater than the 25% percentile and below 1.5 times the mean depth, approximately. Finally, only biallelic sites were retained using BCFtools view with the ‘-M 2’ flags. For specific filters and values see Github https://github.com/ellieearmstrong/Gray_Fox_2023/tree/main/filtering.

#### Sample selection

We performed a principal component analysis (PCA) on SNP genotype data including all 41 individuals using the SNPRelate v1.36.1[^13^](https://app.readcube.com/library/bc53ac12-924a-4977-b177-875ccbf08ca1/all?uuid=38477047084293925&item_ids=bc53ac12-924a-4977-b177-875ccbf08ca1:956038da-ad21-45c5-bd91-b4c81147c149) package in R v4.3.2[^9^](https://app.readcube.com/library/bc53ac12-924a-4977-b177-875ccbf08ca1/all?uuid=778732942041096&item_ids=bc53ac12-924a-4977-b177-875ccbf08ca1:6d003005-8cc7-4751-b838-df235f4833d9). We first converted the VCF file to a genomic data structure file using the snpgdsVCF2GDS function. PCA was conducted with the snpgdsPCA function on genotype data with no missing values, generating eigenvectors for each sample and eigenvalues representing the variance explained by each principal component. Based on the PCA results (Figure S1), we subsampled the dataset to exclude hybrids, which are known to occur in the southwest sampling sites. To achieve this, we selected the westernmost samples in PC space, prioritizing the inclusion of the high-coverage sample, and matched this selection by choosing the same number of easternmost samples.

#### Genome annotations

We produced mappability scores for each genome using GenMap v1.3.0[^14^](https://app.readcube.com/library/bc53ac12-924a-4977-b177-875ccbf08ca1/all?uuid=5054164758307396&item_ids=bc53ac12-924a-4977-b177-875ccbf08ca1:7f8f6878-1f79-41fa-9160-186d8e7a4a84). Mappability scores are used to assess the uniqueness of kmers in the genome and identify regions which are repetitive and cause errors during mapping and variant calling. We used the filterGM.rb script from Armstrong & Campana[^15^](https://app.readcube.com/library/bc53ac12-924a-4977-b177-875ccbf08ca1/all?uuid=602399376045917&item_ids=bc53ac12-924a-4977-b177-875ccbf08ca1:a307bf30-d888-4659-8ec5-b339baceb04b) to generate a file of sites with a mappability score < 1.

We also generated repetitive element annotations for each genome. Briefly, we ran TETools v1.7[^16^](https://app.readcube.com/library/bc53ac12-924a-4977-b177-875ccbf08ca1/all?uuid=05283154733352191&item_ids=bc53ac12-924a-4977-b177-875ccbf08ca1:f8a965e4-221b-4791-81c0-062cf28d8b94) on each genome assembly as in Armstrong et al.[^5^](https://app.readcube.com/library/bc53ac12-924a-4977-b177-875ccbf08ca1/all?uuid=6579376902024113&item_ids=bc53ac12-924a-4977-b177-875ccbf08ca1:5ddd40e9-72a1-497b-8efe-aec57a4ebcd8). Final output files were converted to bed files using the RM2bed.rb script and subsequently combined with the mappability scores file using the simplify_bed.rb script[^15^](https://app.readcube.com/library/bc53ac12-924a-4977-b177-875ccbf08ca1/all?uuid=8432552747290996&item_ids=bc53ac12-924a-4977-b177-875ccbf08ca1:a307bf30-d888-4659-8ec5-b339baceb04b).

Genome annotations were previously generated for both versions of the dog genome, however, annotations have not been previously generated for the gray fox. Ideally, a combination of evidence from RNA and closely related species annotations would be used to annotate the gray fox genome, but RNA data has not been generated for this species. As such, we used the GALBA v1.0.9[^17^](https://app.readcube.com/library/bc53ac12-924a-4977-b177-875ccbf08ca1/all?uuid=0353521628249841&item_ids=bc53ac12-924a-4977-b177-875ccbf08ca1:14bfc7ce-6b5a-4f9f-b4b5-8b6ef413dfd5) program to generate draft annotations for the gray fox. Though these annotations are likely imperfect, since we are only using annotation information to remove regions which may be evolving non-neutrally, these are sufficient for our purposes of exclusion. We downloaded protein files from the arctic fox (GCF_018345385.1[^3^](https://app.readcube.com/library/bc53ac12-924a-4977-b177-875ccbf08ca1/all?uuid=36209445108985805&item_ids=bc53ac12-924a-4977-b177-875ccbf08ca1:d6c8cdc2-b55c-4910-9cde-bc2b0d508642)), domestic dog (GCF_011100685.1[^4^](https://app.readcube.com/library/bc53ac12-924a-4977-b177-875ccbf08ca1/all?uuid=7772257623744803&item_ids=bc53ac12-924a-4977-b177-875ccbf08ca1:a10e08eb-cd9b-4502-ae02-d5757a7bfff3)), brown bear (GCF_023065955.2[^18^](https://app.readcube.com/library/bc53ac12-924a-4977-b177-875ccbf08ca1/all?uuid=49879056844254444&item_ids=bc53ac12-924a-4977-b177-875ccbf08ca1:80cfdc09-e41f-488c-bc1c-bd83482551bd)), and the most recent (at the time) human genome assembly (GCF_009914755.1[^19^](https://app.readcube.com/library/bc53ac12-924a-4977-b177-875ccbf08ca1/all?uuid=48678391637753293&item_ids=bc53ac12-924a-4977-b177-875ccbf08ca1:a72657fd-49f9-412b-8593-599616592a96)). These files were provided as protein evidence to GALBA using the –prot_seq flag and run using AUGUSTUS v3.3.2[^20^](https://app.readcube.com/library/bc53ac12-924a-4977-b177-875ccbf08ca1/all?uuid=5532140738629825&item_ids=bc53ac12-924a-4977-b177-875ccbf08ca1:15f5aafd-756a-48ad-990d-f739394c685e) which was run with default parameters.

#### Functional enrichment analysis of unmapped reads

To determine whether the reads that did not map to heterospecific reference genomes but mapped to the conspecific genome were enriched for functional regions, we used the read tags (unique read identifiers) of the unmapped reads from Arctic fox and CanFam4 genomes to obtain the positions where they mapped in the gray fox genome. We used bedtools v2.31.1[^21^](https://app.readcube.com/library/bc53ac12-924a-4977-b177-875ccbf08ca1/all?uuid=8705483638537324&item_ids=bc53ac12-924a-4977-b177-875ccbf08ca1:dd74bc15-5d33-4a18-98a6-9d9e786b7a1a) to intersect those positions with the gray fox genome annotation, then conducted a functional enrichment analysis for gene ontology biological processes using Enrichr[^22^](https://app.readcube.com/library/bc53ac12-924a-4977-b177-875ccbf08ca1/all?uuid=5122634767470065&item_ids=bc53ac12-924a-4977-b177-875ccbf08ca1:72e37431-c2c0-4fdd-a8b2-15db592dd294) to identify biological processes significantly overrepresented in those regions.

#### Computing genetic diversity (π) and differentiation (F_ST_)

Pairwise genetic diversity (π) within each population and genetic differentiation (*F*_ST_) between populations were computed using VCFtools[^23^](https://app.readcube.com/library/bc53ac12-924a-4977-b177-875ccbf08ca1/all?uuid=36626086585739404&item_ids=bc53ac12-924a-4977-b177-875ccbf08ca1:42bb6f4b-ea1f-43e8-85b9-32e0b25d0c01) v1.14, including six individuals from the east and six individuals from the west. We calculated π using a gVCF file containing all genome sites—both variant and invariant—and kept only sites without missing data for each population. *F*_ST_ was calculated using a VCF file with only variant sites, retaining sites with no missing data across populations. We computed both summary statistics in 50 kb windows. We examined the relationship between genetic diversity and differentiation for each population and reference genome using a Kendall's rank correlation test, given the exponential-like distribution of nucleotide diversity. Additionally, to compare the number of segregating sites per site with nucleotide diversity, we calculated Tajima’s D using VCFtools v1.14 in non-overlapping 50 kb windows based on variant sites for each population and reference genome.

#### Quantifying genetic variation

We used VCFtools[^23^](https://app.readcube.com/library/bc53ac12-924a-4977-b177-875ccbf08ca1/all?uuid=8853723574225266&item_ids=bc53ac12-924a-4977-b177-875ccbf08ca1:42bb6f4b-ea1f-43e8-85b9-32e0b25d0c01) to obtain raw allele counts at all biallelic variant sites, which we summarized to get the total number of variants as well as the number of singletons, i.e., rare variants where an allele is only found once. To calculate an SFS for each population based on each reference genome, we used VCFtools to obtain allele counts at all fully covered (i.e., no missing data) biallelic sites, which we summarized to get a folded SFS. Data summaries were performed using the tidyverse package[^24^](https://app.readcube.com/library/bc53ac12-924a-4977-b177-875ccbf08ca1/all?uuid=07135550123240042&item_ids=bc53ac12-924a-4977-b177-875ccbf08ca1:e827811d-d114-409b-b549-c93619f601cd) in R v. 4.3.2[^9^](https://app.readcube.com/library/bc53ac12-924a-4977-b177-875ccbf08ca1/all?uuid=9236499147727228&item_ids=bc53ac12-924a-4977-b177-875ccbf08ca1:6d003005-8cc7-4751-b838-df235f4833d9). Site frequency spectra were generated using 6 individuals from the east and 6 individuals from the west for the conspecific and heterospecific reference genomes.

SNPs with missing data were removed from these analyses. There were 3,496,241 SNPs in the East and 2,669,112 SNPs in the West out of the 26,180,875 total SNPs that were removed due to missing data with Gray fox reference genome; 11,031,507 SNPs in the East and 6,846,230 SNPs in the West out of the 43,583,377 total SNPs that were removed due to missing data with CanFam4 reference genome; and 11,526,220 SNPs in the East and 7,176,780 SNPs in the West out of the 45,042,040 total SNPs that were removed due to missing data with Arctic fox reference genome.

#### Demographic inference

Demography was inferred using 6 individuals from the east and 6 individuals from the west with smc++[^25^](https://app.readcube.com/library/bc53ac12-924a-4977-b177-875ccbf08ca1/all?uuid=037779311608764354&item_ids=bc53ac12-924a-4977-b177-875ccbf08ca1:101e73f2-e09f-43ee-b70c-18dadd41aa3d) v1.15.4, MSMC2[^26^](https://app.readcube.com/library/bc53ac12-924a-4977-b177-875ccbf08ca1/all?uuid=9742900587595915&item_ids=bc53ac12-924a-4977-b177-875ccbf08ca1:3440195e-9159-489a-818e-11f9171e9769) v2.1.4, and Stairway Plot 2 v2.1.2[^27^](https://app.readcube.com/library/bc53ac12-924a-4977-b177-875ccbf08ca1/all?uuid=21236124096876552&item_ids=bc53ac12-924a-4977-b177-875ccbf08ca1:495fa17c-9267-4c88-91bb-3c7d8887fcce). Smc++ combines the SFS with linkage disequilibrium (LD) information in coalescent hidden Markov models (HMM), MSMC2 uses a coalescent HMM without the SFS, and Stairway Plot 2 relies solely on the SFS.
For smc++, we parsed VCFs to generate input files for each autosomal chromosome using the vcf2smc command in smc++. Because smc++ is not able to distinguish regions of missing data from very long runs of homozygosity, positions of regions with highly repetitive sequence and/or low mappability that were masked prior to variant calling were provided in a bed file and marked as missing data using the -m option in the smc++ data sets. Positions of genic regions were obtained from annotations and similarly masked such that demographies were inferred using only putatively neutral regions of the genome. Further, smc++ calculates the SFS by using a single “distinguished individual” selected from the pool of samples against which all other samples are compared, which may impact demographic inference depending on the choice of distinguished individual. We therefore created multiple input files by varying the identity of the distinguished individual and treating the remaining samples from each population as undistinguished. Input files were combined to generate a composite likelihood estimate by running the estimate command for fitting a population size history to the data six separate times, once for each combination of population and reference genome. Runs were conducted assuming a per-site per-generation mutation rate of 4.5e−9[^28^](https://app.readcube.com/library/bc53ac12-924a-4977-b177-875ccbf08ca1/all?uuid=4083414914906398&item_ids=bc53ac12-924a-4977-b177-875ccbf08ca1:ce56edd1-7547-4290-a0c0-bb6d77cee52a) and 25 EM iterations, with estimates restricted between 2,500 and 500,000 generations since present. A thinning parameter of 1,792, calculated as 1,000×log(6) for the six individuals in our analysis, was used as recommended in the smc++ user guide to control the frequency of conditional SFS emission and incorporate information from the undistinguished portion of the sample. A spline representation of population size history was fit using 12 knots to allow for sufficient flexibility while avoiding over-smoothing. We used a generation time of two years[^29^](https://app.readcube.com/library/bc53ac12-924a-4977-b177-875ccbf08ca1/all?uuid=16271870207446493&item_ids=bc53ac12-924a-4977-b177-875ccbf08ca1:a0ddc7fc-ba2f-4d65-81f3-ad7c5dc67dfd) to convert the output from coalescent units to units of time.

For Stairway Plot 2, we used the folded site frequency spectra that were generated for each population and reference genome above as input. We used all bins of the SFS, total callable sites incorporated the monomorphic sites, a per-site per-generation mutation rate of 4.5e−9[^28^](https://app.readcube.com/library/bc53ac12-924a-4977-b177-875ccbf08ca1/all?uuid=787806141448642&item_ids=bc53ac12-924a-4977-b177-875ccbf08ca1:ce56edd1-7547-4290-a0c0-bb6d77cee52a), a generation time of two years[^29^](https://app.readcube.com/library/bc53ac12-924a-4977-b177-875ccbf08ca1/all?uuid=07558987059250655&item_ids=bc53ac12-924a-4977-b177-875ccbf08ca1:a0ddc7fc-ba2f-4d65-81f3-ad7c5dc67dfd), and 67% of the data was used for training. Stairway Plot 2 also implements a check for over or underfitting using various breakpoints. The breakpoints tested were (2, 5, 7, 10) and the best breakpoint value was selected as one that minimized the log-likelihood.

For MSMC2, we first used BCFtools to remove sites with missing data from the gVCF file for each population and reference genome, generated a BED file of sufficiently covered sites for masking, then split files to generate a mask for each chromosome and a gVCF for each sample and chromosome. We then generated input files using generate_multihetsep.py in the MSMC-tools repository, including a negative mask to remove regions with low mappability scores. We estimated population size histories by specifying pair indices (-I) for each individual, to avoid pairs of haplotypes from different individuals, as recommended for unphased genomes.

To evaluate whether demographic reconstructions improve when using SNPs that successfully lifted over from the heterospecific genomes to the gray fox genome, we re-ran smc++ after masking variants in the Arctic fox and dog genomes that failed to liftover. We then compared these masked demographic trajectories to the original reconstructions.

#### Recombination rate estimation

Recombination rates were estimated using pyrho v0.1.0[^30^](https://app.readcube.com/library/bc53ac12-924a-4977-b177-875ccbf08ca1/all?uuid=305460665540669&item_ids=bc53ac12-924a-4977-b177-875ccbf08ca1:327b7308-680f-4659-bafb-ac6cce75b60d), which takes in the demography from smc++ and linkage information from unphased data to estimate population-specific fine-scale recombination rates per generation[^30^](https://app.readcube.com/library/bc53ac12-924a-4977-b177-875ccbf08ca1/all?uuid=8706195235874694&item_ids=bc53ac12-924a-4977-b177-875ccbf08ca1:327b7308-680f-4659-bafb-ac6cce75b60d). Using pyrho, we first precomputed likelihood tables under the demographic models we inferred from smc++ to account for population size fluctuations across time and the mutation rate (4.5e−9[^28^](https://app.readcube.com/library/bc53ac12-924a-4977-b177-875ccbf08ca1/all?uuid=7717014038534753&item_ids=bc53ac12-924a-4977-b177-875ccbf08ca1:ce56edd1-7547-4290-a0c0-bb6d77cee52a)). We then tested multiple sets of hyperparameters to produce reasonable data from the optimization function. Parameters for block penalty, which controls the smoothness of the resulting map, and window size, the amount of bp before a SNP is ignored, were selected based on their minimization of log(*l_2_*). Block penalty parameters tested were (25, 50, 100, 150, 200, 250) and window sizes tested were (25, 50, 100, 150). After hyperparameter selection (Table 2), we averaged the per-base recombination rate inferred by pyrho into 50 kb windows and converted the values to units of centi-Morgan per megabase by multiplying by 1e8 (100 cM per expected crossover multiplied by 1e6 bases per mb). To compare recombination hotspots across references, we converted window positions between each heterospecific reference and the gray fox reference (described below). Hotspots were defined as windows with recombination rates exceeding two standard deviations above the mean for each reference genome and categorized based on whether they were shared across references or unique to a specific reference.

#### Identifying F_ST_ outliers and conducting functional enrichment analyses

Outliers were defined as windows with F_ST_ values exceeding three standard deviations above the mean for each reference genome. To compare outlier regions across references, we converted window positions between each heterospecific reference and the gray fox reference (described below). We also extended this analysis to compare F_ST_ outlier detection between the two heterospecific genomes. Outliers were categorized based on whether they were shared across references or unique to a specific reference. To identify protein-coding genes within the outlier windows, we used Arctic fox and Canfam4 gene annotations, and the draft annotations we generated for the gray fox (described above). We used the GenomicRanges[^31^](https://app.readcube.com/library/bc53ac12-924a-4977-b177-875ccbf08ca1/all?uuid=6987436264020875&item_ids=bc53ac12-924a-4977-b177-875ccbf08ca1:d109cbf4-4c42-4acb-9062-edccabb2a785) v1.54.1 package in R v. 4.3.2[^9^](https://app.readcube.com/library/bc53ac12-924a-4977-b177-875ccbf08ca1/all?uuid=9764130540109973&item_ids=bc53ac12-924a-4977-b177-875ccbf08ca1:6d003005-8cc7-4751-b838-df235f4833d9) to determine which genes fell within the 50 kb windows of each reference genome. Foreground genes were defined as those located within the outlier windows, while background genes included those located outside the outlier windows. We then conducted a functional enrichment analysis for gene ontology (GO) biological processes using Enrichr[^22^](https://app.readcube.com/library/bc53ac12-924a-4977-b177-875ccbf08ca1/all?uuid=9941357377089541&item_ids=bc53ac12-924a-4977-b177-875ccbf08ca1:72e37431-c2c0-4fdd-a8b2-15db592dd294) to identify processes significantly overrepresented in the foreground gene set compared to the background set. We used the ComplexUpset[^32^](https://app.readcube.com/library/bc53ac12-924a-4977-b177-875ccbf08ca1/all?uuid=10206093360837432&item_ids=bc53ac12-924a-4977-b177-875ccbf08ca1:86312feb-47d6-4ced-85cb-cd172b69a29d) v1.3.3 R package to create upset plots showing the number of identified genes from each reference and their intersection. To investigate the impact of reference genome choice on F_ST_ outlier detection, we examined the specific genes located within unique outlier windows for each heterospecific reference genome. This analysis aimed to determine whether biologically plausible interpretations could be derived from potentially reference-biased regions.

To assess the impact of reference choice on outlier classification, we examined multiple genomic features within matched outlier windows, including evolutionary constraint scores, gene annotations, and the distribution of synonymous and non-synonymous mutations. We obtained PhyloP constraint scores for CanFam4 from the International Dog10K project[^33^](https://app.readcube.com/library/bc53ac12-924a-4977-b177-875ccbf08ca1/all?uuid=9371947969618644&item_ids=bc53ac12-924a-4977-b177-875ccbf08ca1:e89b8b6c-0d1c-4b74-8883-3c03e4dbf1b5). Using bigWigAverageOverBed[^34^](https://app.readcube.com/library/bc53ac12-924a-4977-b177-875ccbf08ca1/all?uuid=8212731802783693&item_ids=bc53ac12-924a-4977-b177-875ccbf08ca1:eb6cc9cf-ae08-4361-ae72-97ba5bb5eb28), we calculated the mean PhyloP score for each 50kb window, matching the F_ST_ windows that we lifted over to align across reference genomes using the GenomicRanges package in R. Additionally, we evaluated the genic and intergenic composition of outlier windows based on annotations from each reference genome. Next, we used SnpEff[^35^](https://app.readcube.com/library/bc53ac12-924a-4977-b177-875ccbf08ca1/all?uuid=4961440383933836&item_ids=bc53ac12-924a-4977-b177-875ccbf08ca1:9d59cdcf-c6fe-4958-9e81-5156471bf5b7) to assess the functional impact of variants within outlier windows by building custom annotation databases for each reference genome using its corresponding gene annotations. We annotated variants in the VCF files separately for each reference, classifying mutations as synonymous or non-synonymous based on their predicted coding effects.

#### Converting coordinates between reference genomes

We performed whole genome alignments between the two heterospecific genomes and the gray fox genome using minimap2[^36^](https://app.readcube.com/library/bc53ac12-924a-4977-b177-875ccbf08ca1/all?uuid=6035541547469926&item_ids=bc53ac12-924a-4977-b177-875ccbf08ca1:930d483d-11f6-4668-9d27-b0d6c5bb3906) v2.14 with parameters --cs -x asm20. We converted the pairwise alignment format computed by minimap2 to a chain file using paf2chain[^37^](https://app.readcube.com/library/bc53ac12-924a-4977-b177-875ccbf08ca1/all?uuid=26382659902883654&item_ids=bc53ac12-924a-4977-b177-875ccbf08ca1:0fcc162e-cb87-49a6-a00a-90eb07229022), then used the Crossmap[^38^](https://app.readcube.com/library/bc53ac12-924a-4977-b177-875ccbf08ca1/all?uuid=5037576261533051&item_ids=bc53ac12-924a-4977-b177-875ccbf08ca1:1715acff-02fd-4e3c-b7cc-deef1433f7a3) v0.7.0 *bed* command to convert SNP positions and the *region* command, with the default (-r) value of 0.85 for the minimum ratio of bases that must remap, to convert window positions in the heterospecific genomes to their corresponding positions in the gray fox genome.

### Supplementary Text

#### Read mapping and pairing is improved with a conspecific reference genome

We found significant differences in the proportion of reads mapped between the conspecific and heterospecific reference genomes. Approximately 99% of gray fox reads successfully mapped to the Arctic fox and CanFam4 reference genomes, while 99.7% of gray fox reads mapped to the gray fox reference genome (χ²_KW_=33.19, P<0.001, Table 1). Gene ontology (GO) analysis of gray fox genes located in reads that failed to map to heterospecific genomes revealed significant enrichment for terms related to sensory perception and immunity, including olfactory reception (GO:0050911) and MHC complex assembly (GO:0002399). This suggests that using heterospecific reference genomes may systematically exclude important functional regions. Furthermore, we found no significant differences in the estimated depth across individuals between the genomes (χ²_KW_=0.98, P=0.613). Average read depths were 9.34× for the Arctic fox, 8.78× when using Canfam4, and 8.66× when using the gray fox reference, showing an inverse relationship with genome assembly size (2.35 Gb, 2.48 Gb, and 2.66 Gb, respectively). However, when examining only the autosomes, read depths were more consistent (Arctic fox: 9.35×, gray fox: 9.22×, Canfam4: 9.21×), with less variation in autosomal genome sizes (Arctic fox: 2.20 Gb, Canfam4: 2.23 Gb, gray fox: 2.27 Gb). Interestingly, we found a substantial and significant difference in the percentage of properly paired reads across genomes, with 89.4%, 90.3%, and 94.7% of reads properly pairing when mapped to the Arctic fox, Canfam4, and the gray fox reference genome, respectively (χ²_KW_=40.14, P<0.001). On average, almost 5% more reads were properly paired when mapped to the gray fox reference compared to the two heterospecific genomes.

#### Reference genome choice affects recombination hotspot detection

We defined hotspots as windows with recombination rates exceeding two standard deviations above the mean for each reference genome, then matched these windows across reference genomes to identify shared and unique hotspots. The effect of reference genome choice on hotspot detection differed across populations (Figure S4). In the eastern population, the conspecific gray fox reference identified more recombination hotspots than the heterospecific references, with the Arctic fox genome detecting about half as many unique hotspots as the gray fox genome. In contrast, in the western population, the pattern was reversed, with the heterospecific references identifying substantially more hotspots. Both the Arctic fox and CanFam4 genomes detected more than five times the number of unique hotspots compared to the gray fox reference. For example, in the eastern population, we found 804 unique hotspots with the gray fox reference versus 442 with the Arctic fox reference, with only 34 shared between them, while in the western population, the dog genome detected 1,084 unique hotspots compared to just 176 with the gray fox reference, sharing only 9 hotspots (Figure S4). The limited overlap in hotspots across references suggests that reference genome choice has a strong influence on inferred recombination landscapes, with heterospecific references leading to an excess of inferred hotspots in the west but not in the east.

#### Genomic context of F_ST_ outliers depends on reference choice

We compared PhyloP conservation scores across *F*_ST_ windows matched between references to assess whether outlier regions occur in more constrained genomic regions. In both comparisons—gray fox vs. CanFam4 and gray fox vs. Arctic fox—*F*_ST_ outlier windows had higher mean PhyloP scores than non-outliers (χ²_KW_=66, P<0.001). Outliers had a mean PhyloP score of 0.28, while non-outliers had 0.19. In both cases, outliers were significantly more conserved than non-outliers (Wilcoxon pairwise tests, P<0.001), but there was no significant difference in conservation scores between outliers unique to each reference and those shared with gray fox (P>0.05). Furthermore, we classified matched outlier windows as genic or intergenic based on the genome annotations for the three references. Genic regions were consistently overrepresented in outliers across all three genome annotations, with the highest enrichment in windows that were outliers only in the gray fox genome (Figure S7). Intergenic regions were consistently underrepresented, but their relative enrichment varied across genome annotations, with the highest proportions based on the Arctic fox, followed by CanFam4. Using SnpEff, we classified mutations within outlier windows as synonymous or non-synonymous across the three genome annotations. A striking difference emerged between heterospecific and conspecific references in the relative proportions of mutation types (Figure S7). In the Arctic fox and CanFam4 annotations, synonymous mutations were nearly twice as frequent as non-synonymous mutations (2,315 vs. 1,371 in Arctic fox; 2,544 vs. 1,626 in CanFam4). In contrast, when using the gray fox annotation, non-synonymous mutations outnumbered synonymous ones (1,518 vs. 1,379). Despite these differences, the total number of outlier mutations was similar across references (245 in Arctic fox, 248 in CanFam4, and 285 in gray fox), suggesting that the annotation-driven variation in synonymous vs. non-synonymous counts reflects differences in gene models rather than differences in the number of outlier variants detected.

### Supplementary Figures

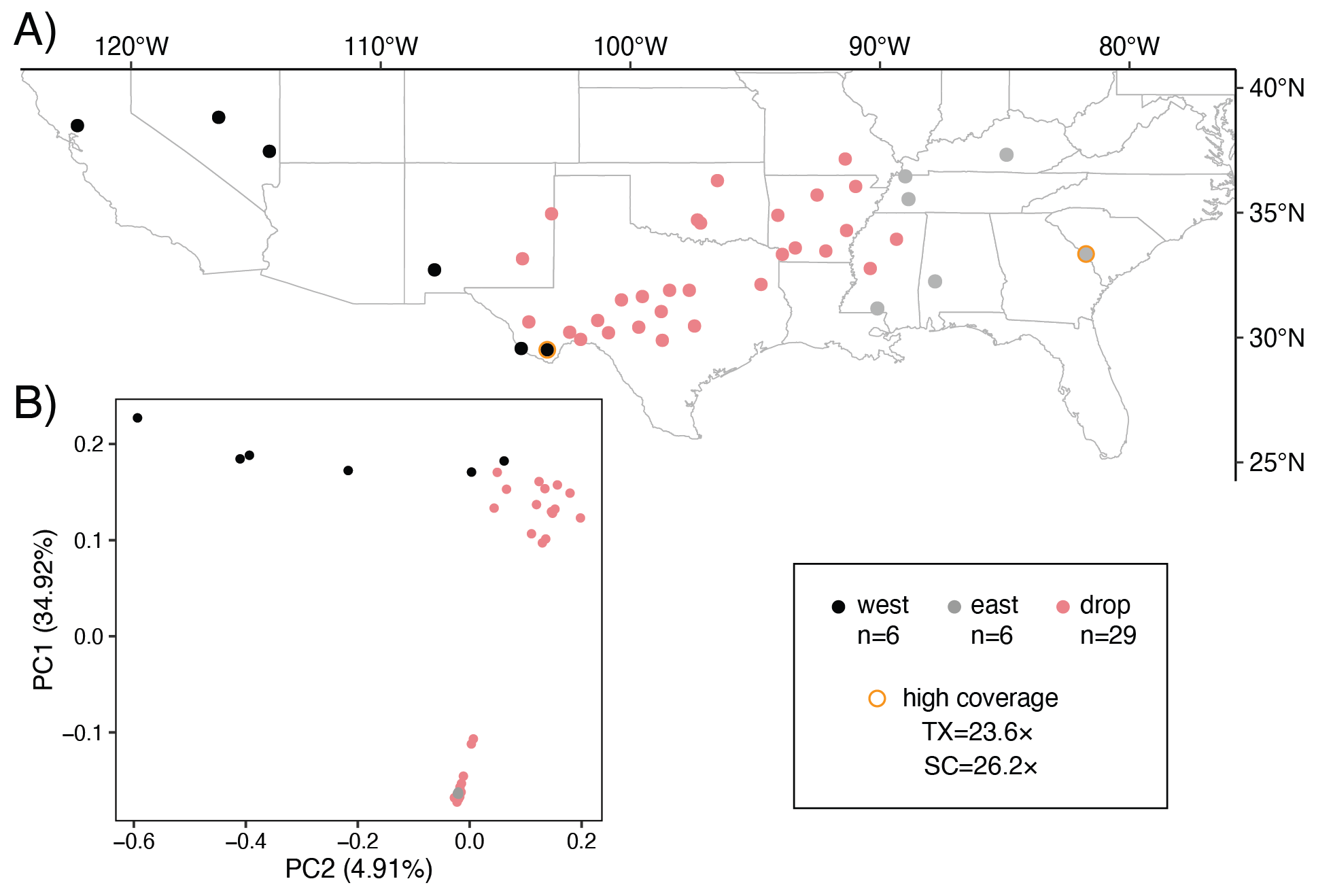

Figure S1. Sampling locations of genomes. A) Sampling localities of WGS data for n=41 gray foxes obtained from Preckler-Quisquater et al.[1](https://app.readcube.com/library/bc53ac12-924a-4977-b177-875ccbf08ca1/all?uuid=46491052237259956&item_ids=bc53ac12-924a-4977-b177-875ccbf08ca1:9d9c5b74-25b4-4d11-b82a-6467769efd3f) with gold borders highlighting high-coverage samples. B) Principal components 1 and 2 of genome-wide variation among individuals. In both panels, black and gray points represent the 12 samples included in our analysis, and the pink points represent the excluded samples, which include admixed individuals.

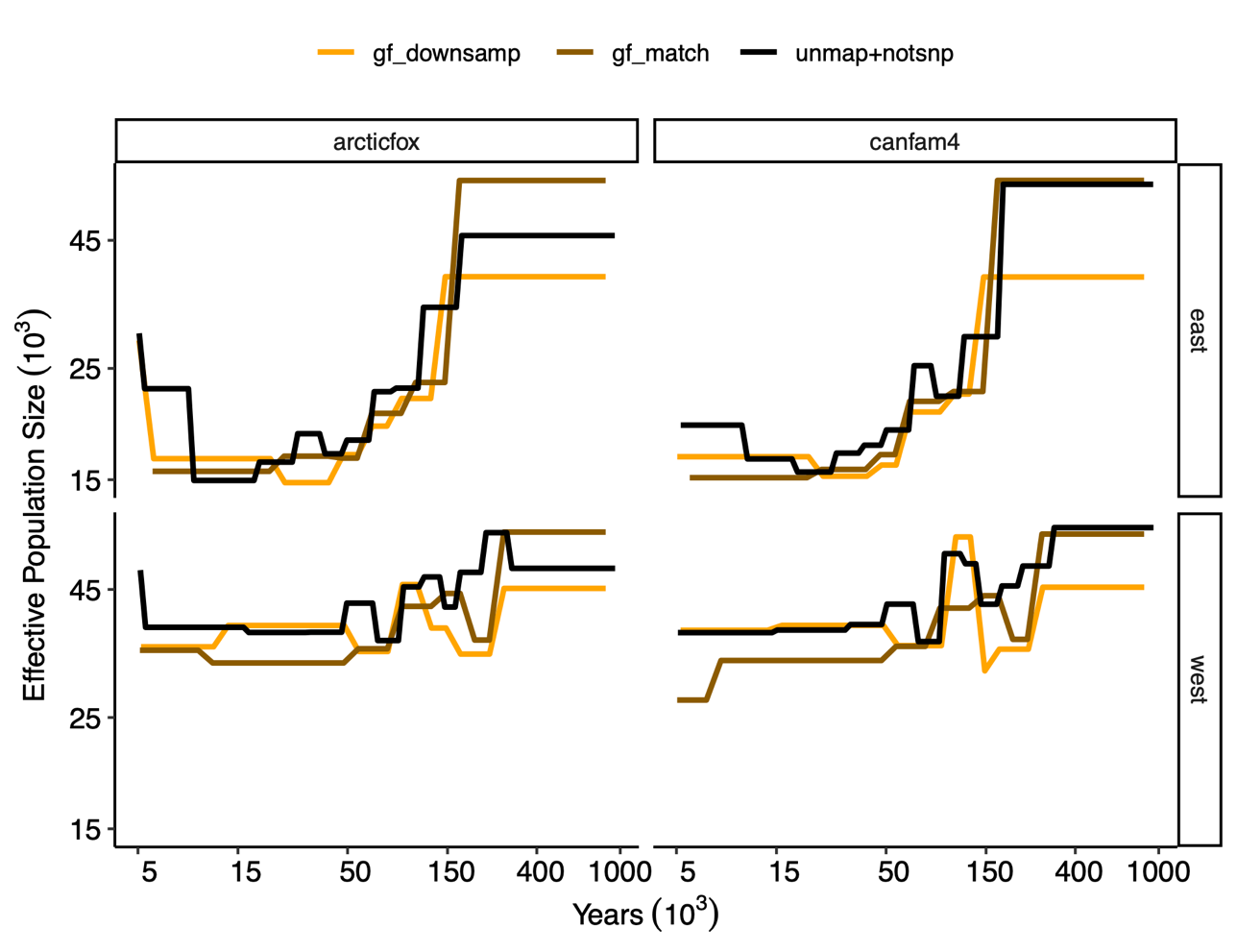

Figure S2. Demographic trajectories using matched SNP sets across reference genomes. Inferred effective population sizes (y-axis) over time in years from present (x-axis) for eastern and western populations using Arctic fox and CanFam4 reference genomes. Black lines: trajectories based on heterospecific genomes using only lifted-over SNPs (invariant sites and unmapped sites masked). Brown lines: trajectories based on gray fox genome using only SNPs that successfully lifted over to the respective heterospecific genome. Gold lines: trajectories based on gray fox genome with randomly downsampled SNPs matching the number of variants that lifted over from each heterospecific genome.

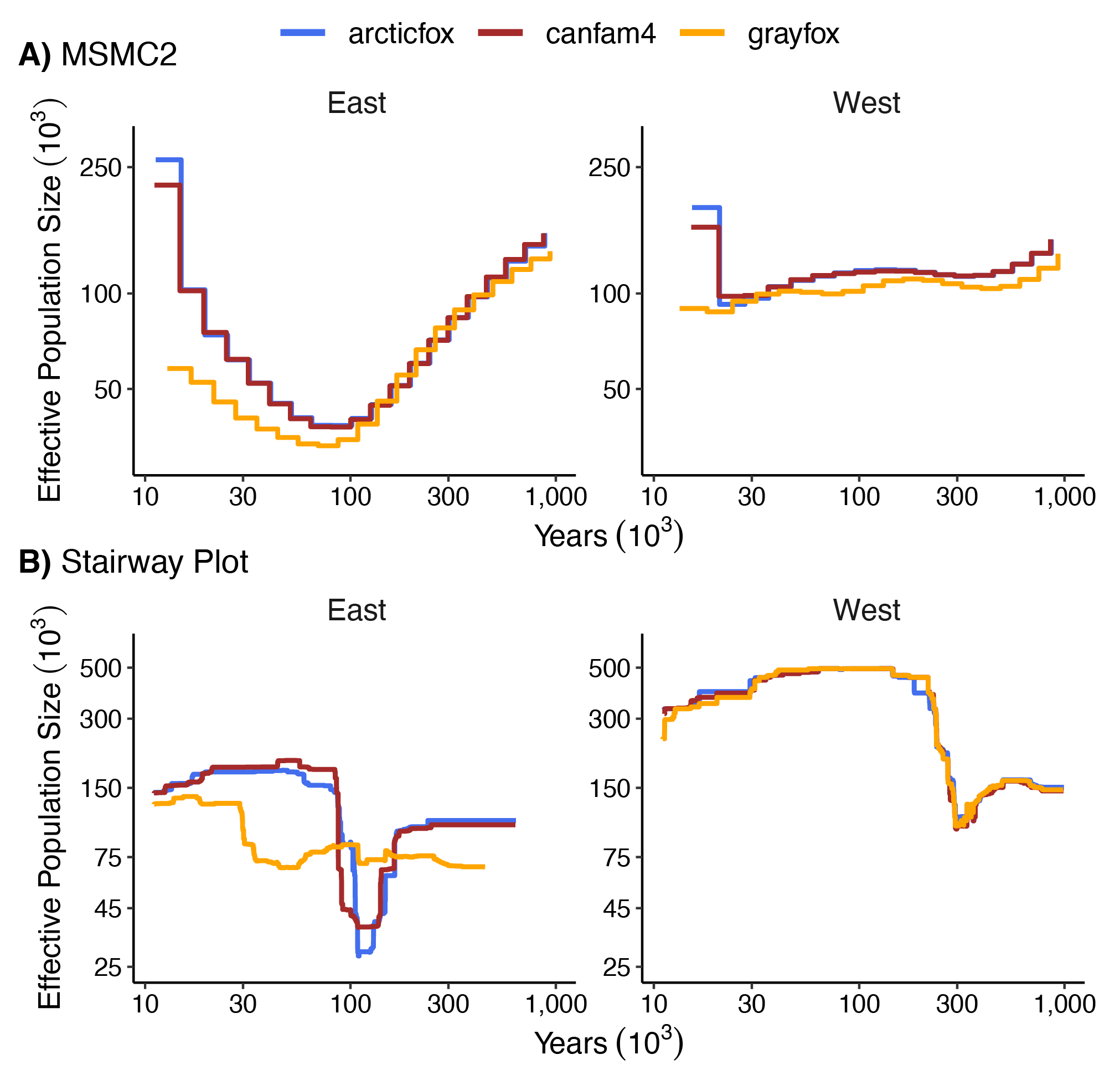

Figure S3. Demographic trajectories across reference genomes. Inferred effective population sizes (y-axis) and years from present (x-axis) reveal discordant demographic histories of foxes resolved using the species-matched (gold) and heterospecific genomes (red and blue) in the east (left) and the west (right) using A) MSMC2 and B) Stairway Plot.

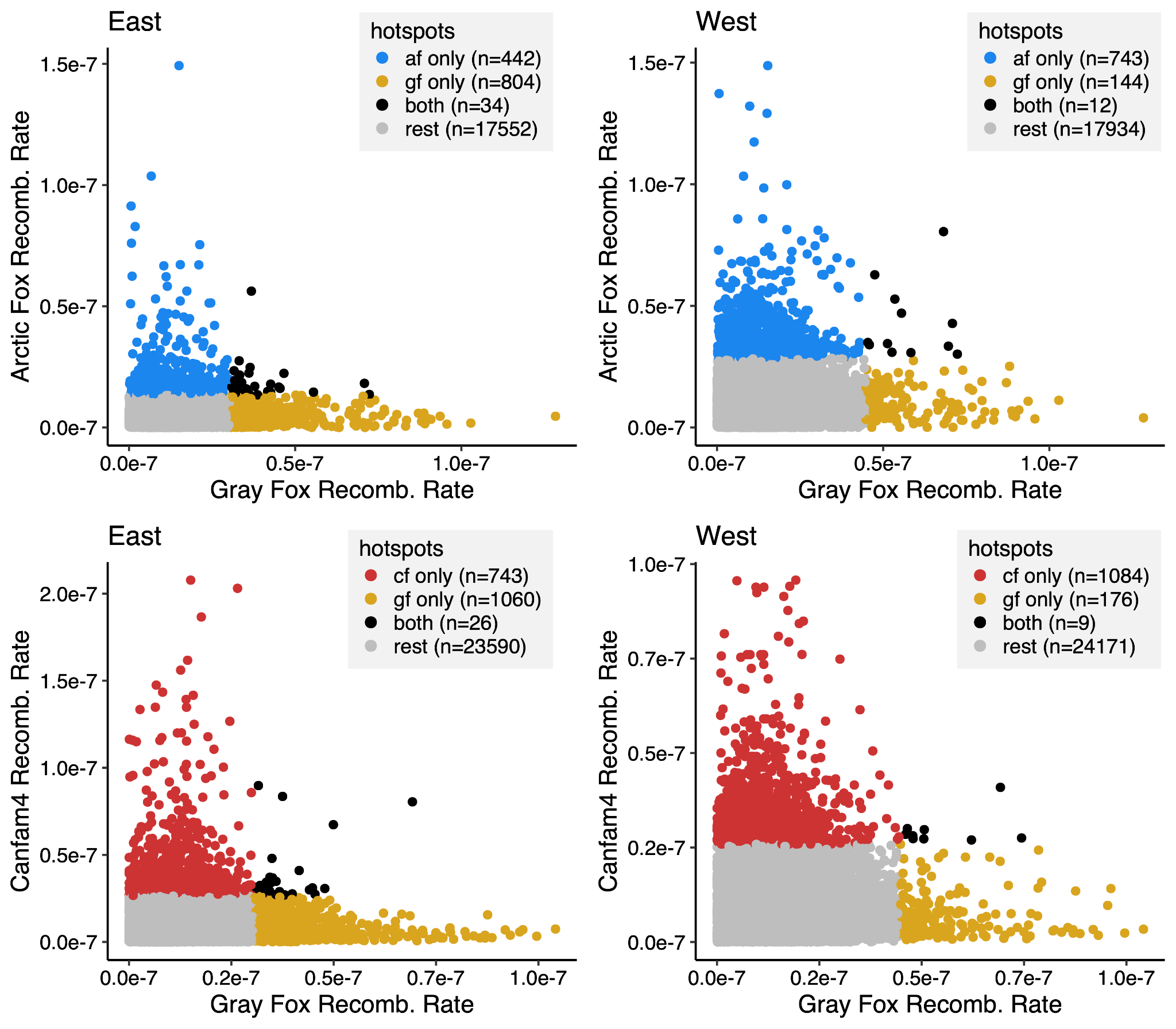

Figure S4. Inferred recombination rates across reference genomes. Scatter plots comparing recombination rates in 50-kb windows between gray fox and heterospecific references for east (left) and west (right) populations. Recombination hotspots unique to the arctic fox (blue, top), Canfam4 (red, bottom) and gray fox (gold) are highlighted. Hotspots shared between the two references are shown in black, and the remaining windows are in gray.

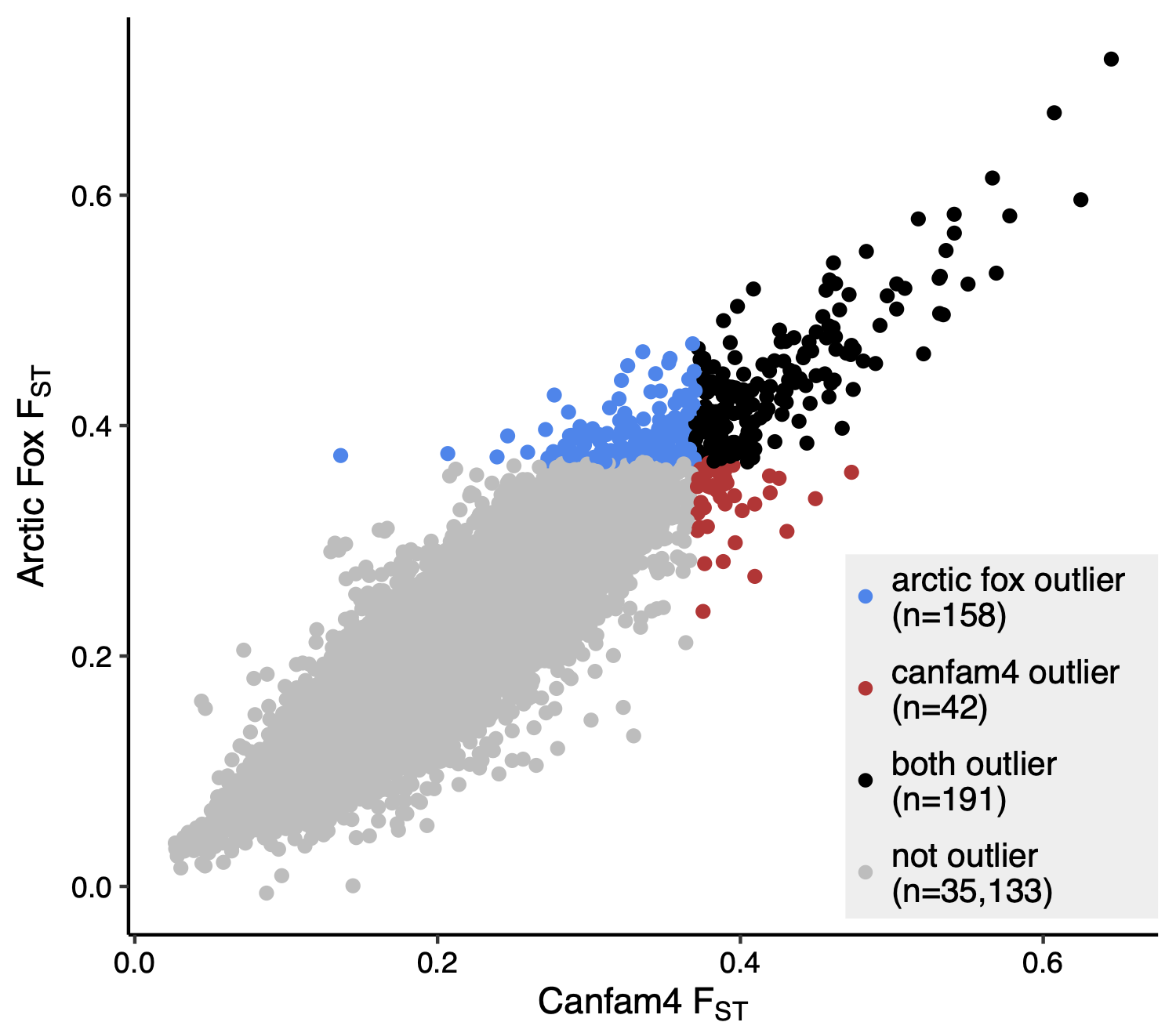

Figure S5. Outlier statistics compared across reference genomes. Scatter plot comparing F_ST_ values between dog (cf4) and arctic fox (af) references, with outlier windows unique to the arctic fox (blue) and Canfam4 (red) highlighted.

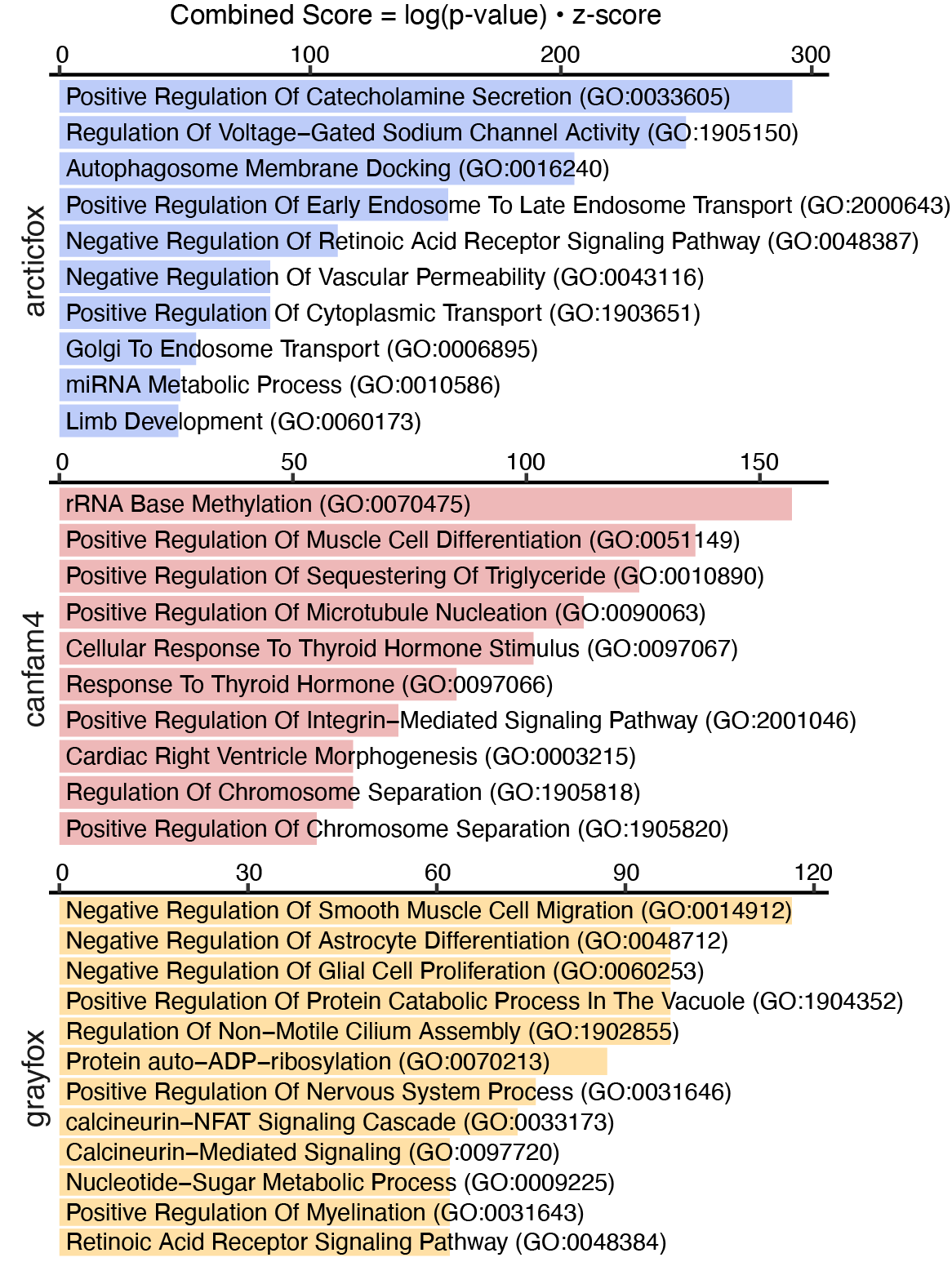

Figure S6. Top 10 unique GO biological process terms identified for each reference genome, highlighting distinct thematic focuses. The unique terms with the highest combined scores are shown for arctic fox (blue), Canfam4 (red), and gray fox (yellow). Each term is listed alongside its GO identifier, with the combined score on the x-axis calculated as the log(p-value) multiplied by the z-score.

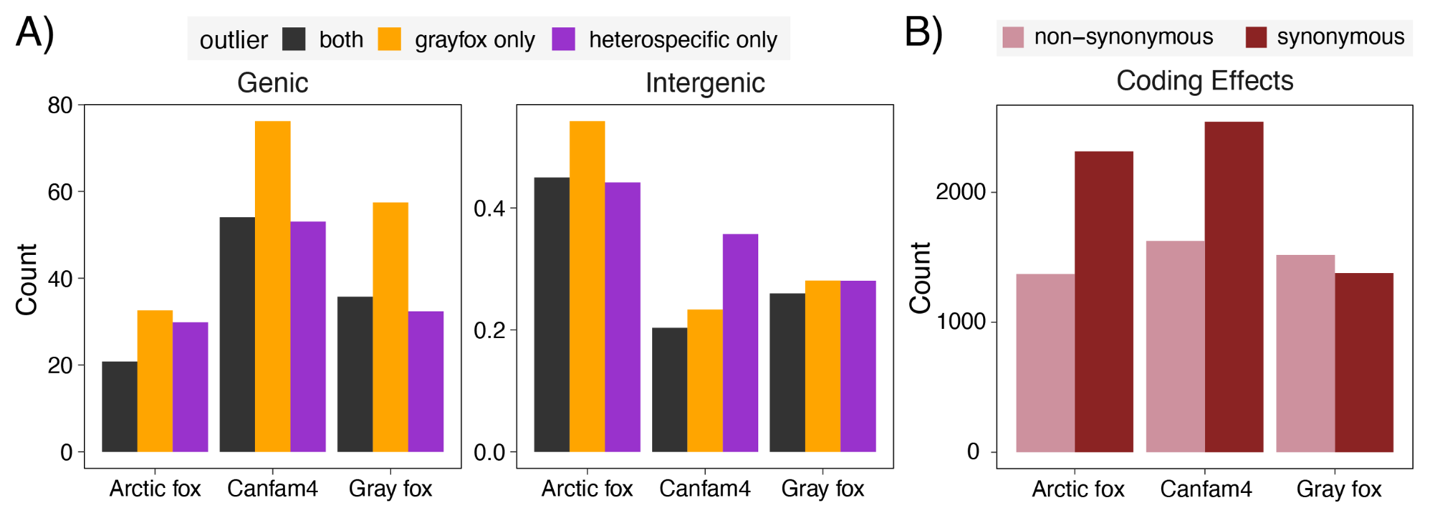

Figure S7. Functional context of F_ST_ outliers by reference genome. (A) Genic (left) and intergenic (right) content of outlier windows based on annotations for the three references, with shared outliers (present in both the heterospecific and conspecific genomes) shown in black, outliers unique to gray fox shown in gold, and outliers unique to Arctic fox or Canfam4 shown in purple. (B) Number of non-synonymous (pink) and synonymous (red) mutations in outlier windows based on each reference.

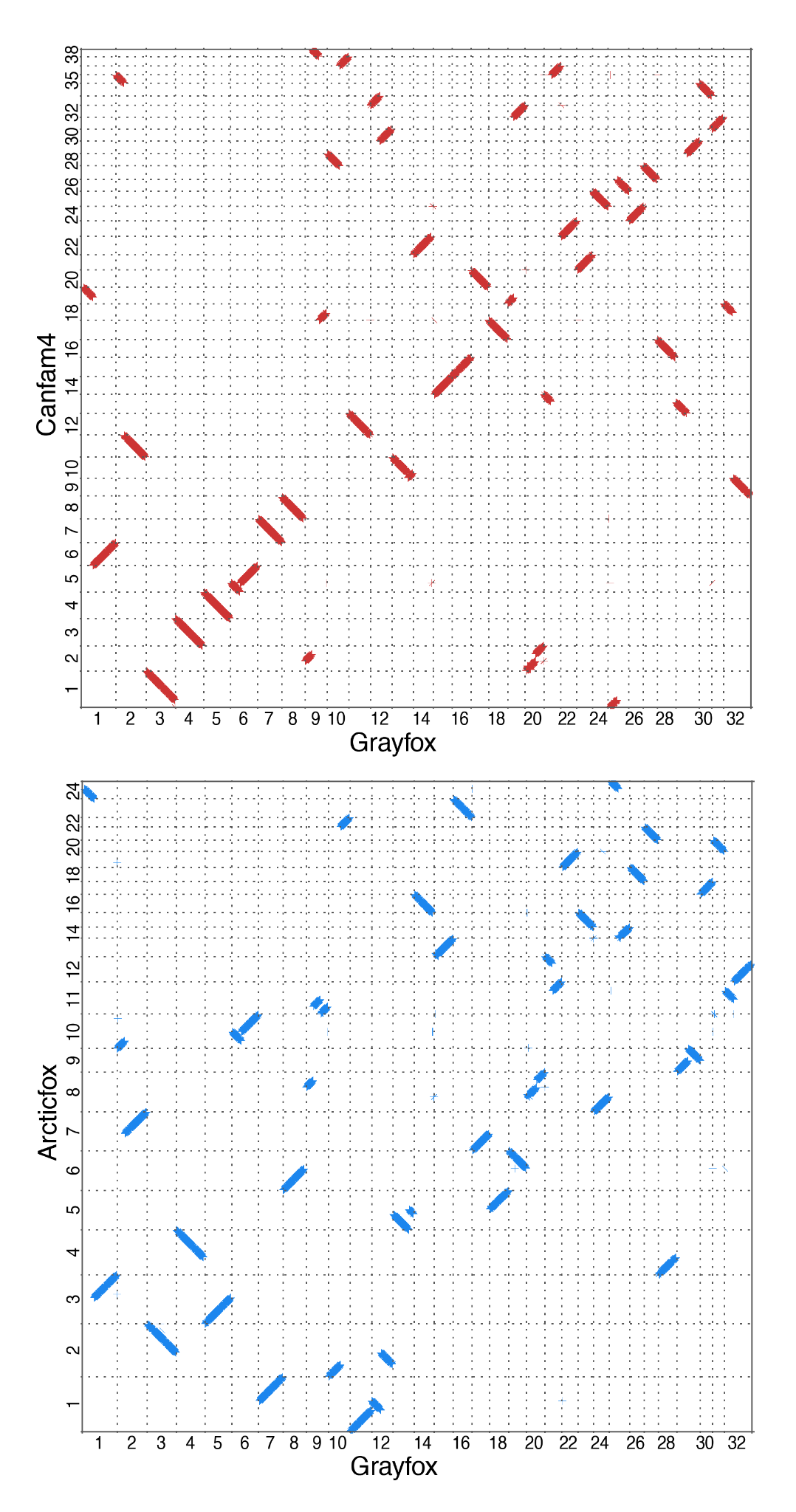

Figure S8. Dot plot of pairwise whole-genome alignment. Plots are between the gray fox genome (x-axis) and the heterospecific reference genomes (y-axis, top) Canfam4 and the Arctic fox (y-axis, bottom) showing synteny is not conserved, with evidence of chromosomal fusions and fissions.

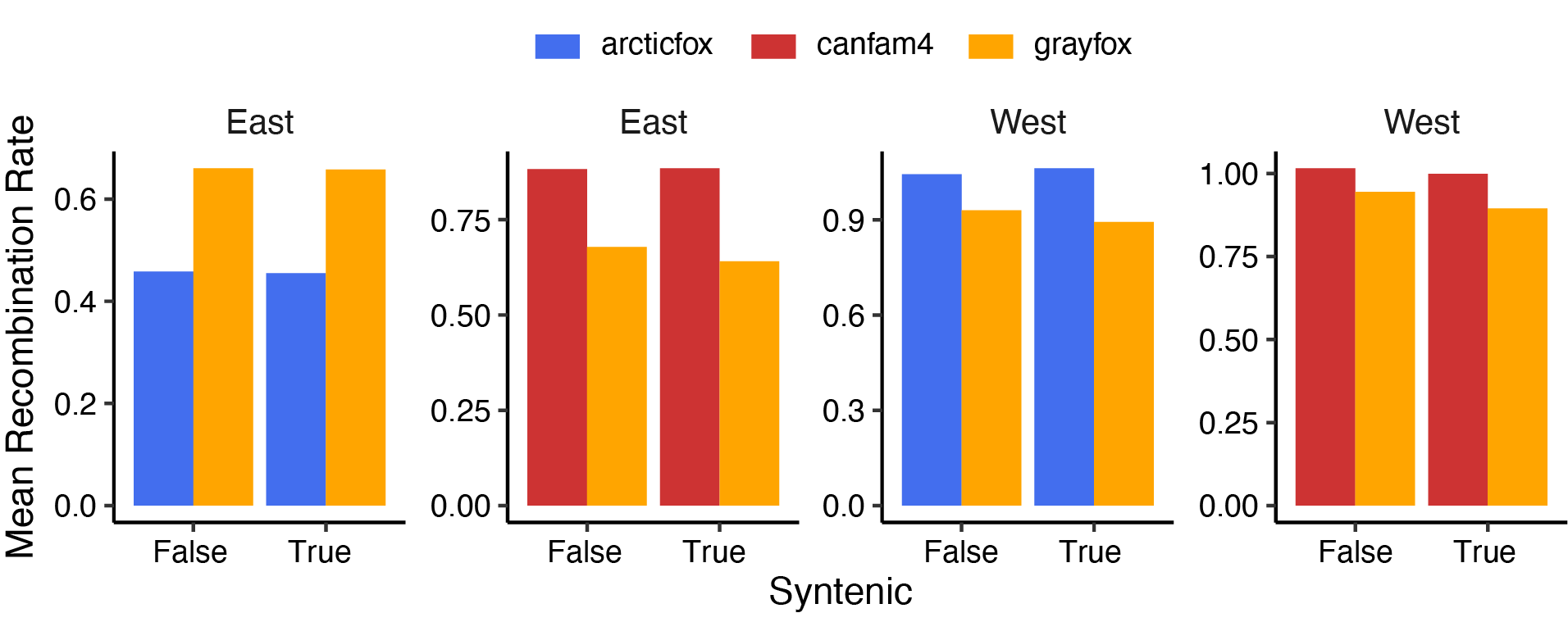

Figure S9. Mean recombination rates for syntenic vs non-syntenic chromosomes between reference genomes. Recombination rates vary more across references, than between syntenic and non-syntenic regions.

### Supplementary Tables

Table S1. Variants from eastern and western gray fox populations that did not lift over from CanFam4 and Arctic fox reference genomes to the gray fox reference genome. Liftover outcomes are classified first as either "not SNP" (meaning the site successfully mapped but is monomorphic in the gray fox genome) or "not map" (meaning the site failed to map to the heterospecific genome). Then, we partition the sites in each category as either singletons or SNPs (meaning non-singletons), in the heterospecific genome for each population. The count represents the absolute number of variants in each category, and percent represents the proportion of the total variants from that population that fell into each category.

| Population | Genome | Gray Fox | Heterospecific SNP | Count | Percent |
| --- | --- | --- | --- | --- | --- |
| East | CanFam4 | Not SNP | Singleton | 311,143 | 5.8% |
|  |  |  | SNP | 254,069 | 4.7% |
|  |  | Not Map | Singleton | 114,321 | 2.1% |
|  |  |  | SNP | 252,029 | 4.7% |
|  | Arctic Fox | Not SNP | Singleton | 297,217 | 5.5% |
|  |  |  | SNP | 257,776 | 4.8% |
|  |  | Not Map | Singleton | 131,316 | 2.4% |
|  |  |  | SNP | 298,266 | 5.5% |
| West | CanFam4 | Not SNP | Singleton | 391,088 | 3.9% |
|  |  |  | SNP | 308,813 | 3.1% |
|  |  | Not Map | Singleton | 241,220 | 2.4% |
|  |  |  | SNP | 402,966 | 4.0% |
|  | Arctic Fox | Not SNP | Singleton | 392,752 | 3.9% |
|  |  |  | SNP | 310,697 | 3.1% |
|  |  | Not Map | Singleton | 282,787 | 2.8% |
|  |  |  | SNP | 475514 | 4.7% |

Table S2. Syntenic chromosomes between gray fox and A) dog and B) Arctic fox genomes.

| A) | Canfam4 | Gray fox | B) | Arctic fox | Gray fox |
| --- | --- | --- | --- | --- | --- |
|  | 3 | 4 |  | 13 | 15 |
|  | 4 | 5 |  | 15 | 23 |
|  | 7 | 7 |  | 16 | 14 |
|  | 8 | 8 |  | 17 | 30 |
|  | 10 | 13 |  | 18 | 26 |
|  | 12 | 11 |  | 19 | 22 |
|  | 14 | 15 |  | 20 | 31 |
|  | 15 | 16 |  | 21 | 27 |
|  | 16 | 28 |  | 23 | 16 |
|  | 17 | 18 |  |  |  |
|  | 20 | 17 |  |  |  |
|  | 21 | 23 |  |  |  |
|  | 24 | 26 |  |  |  |
|  | 27 | 27 |  |  |  |
|  | 31 | 31 |  |  |  |
|  | 34 | 30 |  |  |  |
